# Hierarchical Analysis of Multi-mapping RNA-Seq Reads Improves the Accuracy of Allele-specific Expression

**DOI:** 10.1101/166900

**Authors:** Narayanan Raghupathy, Kwangbom Choi, Matthew J. Vincent, Glen L. Beane, Keith Sheppard, Steven C. Munger, Ron Korstanje, Fernando Pardo-Manual de Villena, Gary A. Churchill

## Abstract

Allele-specific expression (ASE) refers to the differential abundance of the allelic copies of a transcript. Direct RNA sequencing (RNA-Seq) can provide quantitative estimates of ASE for genes with transcribed polymorphisms. However, estimating ASE is challenging due to ambiguities in read alignment. Current approaches do not account for the hierarchy of multiple read alignments to genes, isoforms, and alleles. We have developed EMASE (Expectation-Maximization for Allele Specific Expression), an integrated approach to estimate total gene expression, ASE, and isoform usage based on hierarchical allocation of multi-mapping reads. In simulations, EMASE outperforms standard ASE estimation methods. We apply EMASE to RNA-Seq data from F1 hybrid mice where we observe widespread ASE associated with cis-acting polymorphisms and a small number of parent-of-origin effects at known imprinted genes. The EMASE software is freely available under GNU license at https://github.com/churchill-lab/emase and it can be adapted to other sequencing applications.

## Background

Allele-specific expression (ASE) refers to the relative abundance of the two alleles of a transcript in a diploid organism. ASE can result from differential rates of transcription, differences in mRNA stability, or other mechanisms that affect transcript abundance. Allelic differences can range in magnitude from subtle quantitative effects to purely monoallelic expression. ASE is driven by factors that are linked to the gene and act in *cis* to affect transcript abundance. These *cis*-acting factors may be local genetic variants or epigenetic marks that distinguish maternal and paternal alleles. In the absence of *cis*-acting variation, *trans*-acting factors should exert an equal influence on both allelic copies of a gene. ASE has been associated with genetic predisposition to disease (De La Chapelle, 2009; McKean *et al*, 2016). Accurate estimation of ASE can provide insight into mechanisms of normal transcriptional regulation and it can reveal allelic dysregulation that may underlie or reflect disease states (Wittkopp *et al*, 2004).

Early studies of genome-wide ASE used specialized microarray platforms (Ronald *et al*, 2005) and hybridization artifacts posed significant problems for accurate quantitation (Gresham *et al*, 2010). The advent of massively parallel RNA sequencing technologies (RNA-Seq) has provided a unique opportunity to measure ASE directly (Lister *et al*, 2008; Nagalakshmi *et al*, 2008; Mortazavi *et al*, 2008; Wang *et al*, 2009). But the analysis of ASE from RNA-Seq data presents new challenges. In particular, while transcribed genetic variation provides the information needed to discriminate the allelic origin of a transcript, allelic differences can also introduce systematic biases in alignment and ASE estimates (Degner *et al*, 2009).

Numerous RNA-Seq experiments have been carried out to measure ASE and to evaluate the relative contributions of genetic variation and parental imprinting in humans (Lappalainen *et al*, 2013; Baran *et al*, 2015; Babak *et al*, 2015; Kukurba *et al*, 2014) and in model organisms, including yeast (Skelly *et al*, 2011), flies (McManus *et al*, 2010; Coolon *et al*, 2012; Graze *et al*, 2012), and mice (Gregg *et al*, 2010; DeVeale *et al*, 2012; Goncalves *et al*, 2012; Lagarrigue *et al*, 2013). Estimates of the extent of parent-of-origin effects reported in these studies vary widely. While biological differences among organism, cell type, and developmental stage may account for some of this variability, it is also likely that technical differences in experimental design, analysis methods, and interpretation have contributed to the discordance (DeVeale *et al*, 2012; Coolon *et al*, 2012).

Quantification of ASE from RNA-Seq data begins with alignment of sequence reads to a genome or transcriptome — the set of all transcripts derived from a genome. Polymorphisms between the parental genomes enable the allelic origin of some reads to be unambiguously determined and these reads provide the information needed to estimate ASE. Some of the first attempts to estimate ASE from short read RNA-Seq data aligned reads to a haploid reference genome, allowing for mismatches in the alignment, and then counted allelic proportions at known single nucleotide polymorphisms (SNP) loci (Gregg *et al*, 2010; Andergassen *et al*, 2015). However, it is now recognized that alignment to a reference genome can bias estimation in favor of the allele that is most similar to the reference (Degner *et al*, 2009; Stevenson *et al*, 2013; Munger *et al*, 2014). Several approaches have been developed to reduce this bias by accounting for known SNPs in the scoring of alignments (Montgomery *et al*, 2010; Pickrell *et al*, 2010; Lalonde *et al*, 2011; Rozowsky *et al*, 2011; Stevenson *et al*, 2013; Castel *et al*, 2015). Other methods explicitly represent haploid maternal and paternal genomes that incorporate known genetic variation but perform alignment sequentially (Rozowsky *et al*, 2011). ASE estimation can also be improved by assigning reads to alleles of a transcript based on all known SNPs (Wang *et al*, 2011; Coolon *et al*, 2012). These approaches reduce but do not fully remove biases that arise from the initial reference alignment. For example, van de Geijn et al (2015) showed that reads from non-reference alleles frequently map to multiple genomic locations and would be discarded by these methods. Ideally, all of these challenges — diploid alignment, using information in multiple SNPs, and accounting for ambiguous read alignments — should be addressed in a unified statistical analysis framework.

In this work we focus on alignment to a diploid transcriptome, which includes sequences from both allelic copies of all transcript isoforms. The diploid transcriptome has a natural hierarchical structure. Genes, the transcribed regions of the genome, are present as two copies, the maternal and paternal alleles, either of which can be transcribed and processed into multiple different isoforms. A transcript originates from one isoform of one allele of one gene but different transcript sequences may be highly similar or even identical to one another. As a result, a short read sequence may align equally well to multiple transcript sequences. Alignment ambiguities can occur at different levels of the hierarchy. Sequence similarity shared across genes can give rise to *genomic* multi-reads that align to multiple locations in the genome. Exon or exon-junction sharing between transcripts can result in *isoform* multi-reads that align to more than one isoform of the same gene. Lastly, the absence of distinguishing polymorphisms can give rise to *allelic* multi-reads that align equally well to both allelic copies of a gene. A single read can display multiple ambiguities at different levels of this hierarchy. Accounting for multi-mapping reads is known to improve estimation of transcript abundance but little attention has been given to the role of these different types of multi-reads in the estimation process.

One approach to resolve multi-reads is to employ an expectation maximization (EM) algorithm to assign probabilistic weights that apportion the read across multiple transcripts. Previously reported EM algorithms for RNA-Seq analysis do not differentiate between genomic, isoform, and allelic multi-reads (Li and Dewey, 2011; Nicolae *et al*, 2011; Turro *et al*, 2011; Roberts and Pachter, 2013; Patro *et al*, 2014; Bray *et al*, 2016). Here we report an EM algorithm that accounts for the hierarchical structure of the transcriptome. Our method is implemented in open source software, EMASE (https://github.com/churchill-lab/emase). We describe the EMASE algorithm and evaluate its performance using simulated and real data. We use simulated data to evaluate four EMASE models with different hierarchies and compare the performance of EMASE to several widely used methods for estimating ASE. We demonstrate the application of EMASE to real data by analyzing liver RNA-Seq data from a reciprocal F1 hybrid cross between two inbred mouse strains.

## Results

### Importance of counting multi-reads

RNA-Seq data consist of millions of sequence reads obtained from an RNA sample. We represent the transcriptome as a collection of sequence elements, one for each allele of each isoform of each gene, and we assume that each read originated from exactly one element. Some elements of the transcriptome may be highly similar or even identical to one another. There are sequence similarities across gene families; isoforms of a gene may share exons or exon-junctions; and alleles may have a few or no distinguishing polymorphisms. As a result, a read may align to one or more elements in the transcriptome with equal alignment quality. If the best alignment is unique we assume it is correct. Otherwise, we assume the read originated from exactly one of the elements with equally best alignment quality.

Discarding ambiguous or multi-mapping reads is unfortunately a common practice in RNA-Seq analysis (reviewed in Conesa et al (2016)). In addition to loss of information, selectively discarding reads can bias results. The impact of discarding genomic multi-reads on total gene expression has been documented (Li *et al*, 2010; Robert and Watson, 2015). Relatively less attention has been paid to the impact of discarding isoform and allelic multi-reads but it remains a standard practice to discard these reads on the assumption that they are uninformative (Kanitz *et al*, 2015; Castel *et al*, 2015).

To illustrate the potential impact of multi-reads, we counted the different classes in our F1 cross data (Figure 1). Only ~14% of all aligned reads are unique (U in Figure 1) at all levels and the remaining ~86% of reads are multi-reads for at least one level of the hierarchy. Simple multi-reads are multiply aligned at exactly one level of the hierarchy; they represent 42% of all reads (G+I+A). Complex multi-reads are multiply aligned at 2 or more levels in the hierarchy; they represent 44% of all reads (GI+GA+IA+GIA). Thus complex multi-reads represent a significant fraction of the total data, and information; these are the reads that are apportioned in different ways depending on what we assume about the hierarchy of genes, alleles and isoforms.

Unique reads are simply the complement of multi-reads. They provide critical information needed to assign weights and allocate multi-reads. The majority of reads (83%) are genomic unique (A+I+AI+U). Reads that are both genomic and allelic unique represent 22% (I+U) of the total; these reads are most informative for ASE. In the diploid transcriptome of our F1 animals, 88% of genes have at least one allelic variant site; for genes with no variants there will be no allelic unique reads and we cannot estimate ASE. Reads that are both genomic and isoform unique represent a larger proportion, 48% (A+U) of total reads. However, many of these reads align to single-isoform genes (36% of total reads) and thus only 12% of total reads are informative for distinguishing among isoforms.

**Figure 1:**
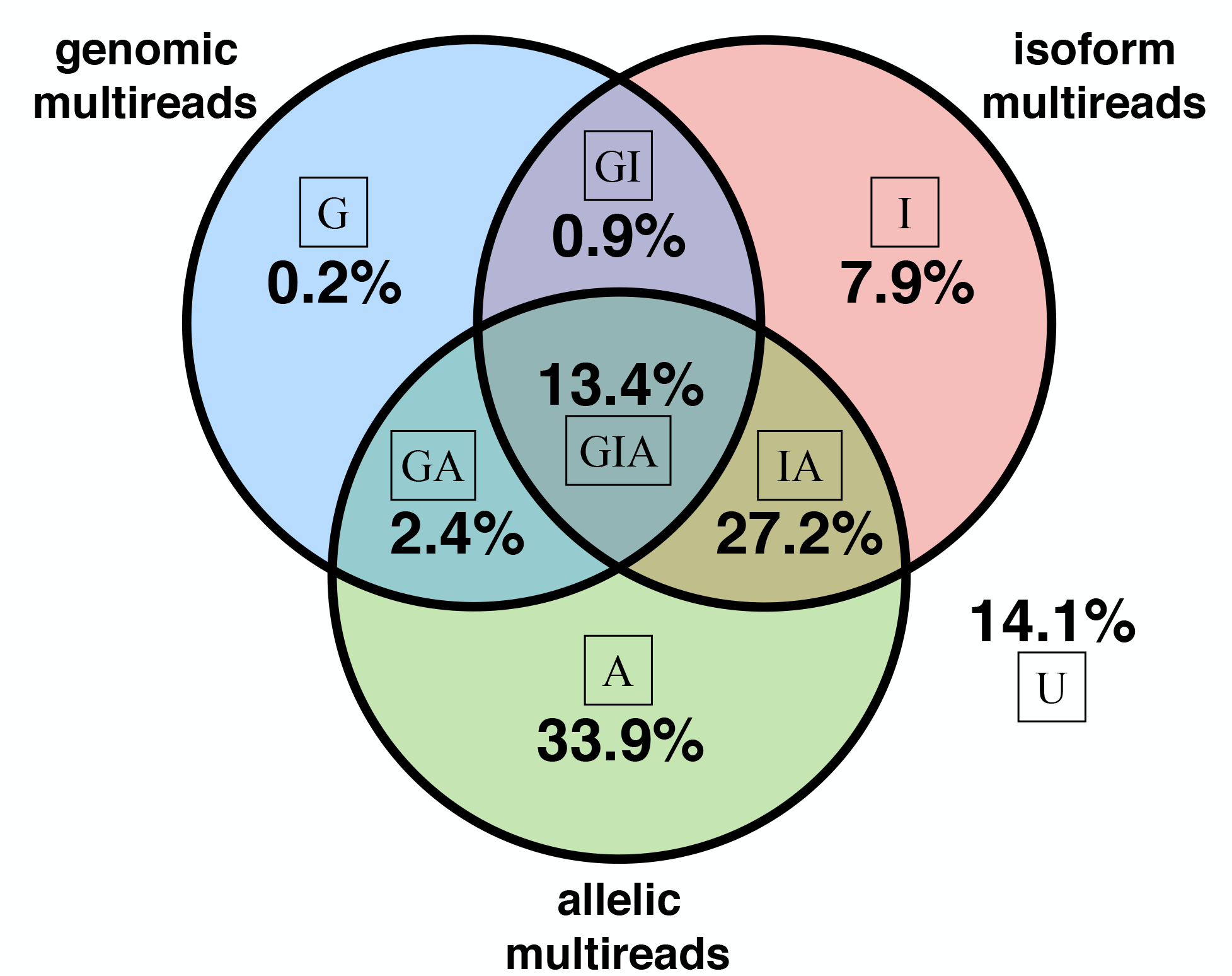
Multi-read proportions in hybrid mouse data. For each read we determined whether it aligns to multiple genomic locations, multiple isoforms of a gene, and multiple alleles. If, for example, a read is a genomic multi-read and is also an isoform multi-read for at least one of its genomic alignments, the read is counted as an isoform multi-read. Complex multi-reads are shown at the intersections of the Venn diagram. The proportion of reads that align uniquely at all levels is 14.1% as shown.

### Fitting an EMASE model

We address the problem of resolving multi-reads in RNA-Seq data by assigning probabilistic weights to each alignment of a multi-read. Current approaches to resolve multi-reads treat all alignments equally regardless of whether the multiple alignments involve alleles, isoforms or genes (Li and Dewey, 2011; Nicolae *et al*, 2011; Turro *et al*, 2011; Roberts and Pachter, 2013; Patro *et al*, 2014; Bray *et al*, 2016). This approach ignores the hierarchical structure of the transcriptome in which a gene may have multiple isoforms and each isoform will have two alleles. As noted above, a single read can be a multi-read at multiple levels in this hierarchy. It is not immediately obvious how to allocate weights for these complex multi-reads.

**Figure 2:**
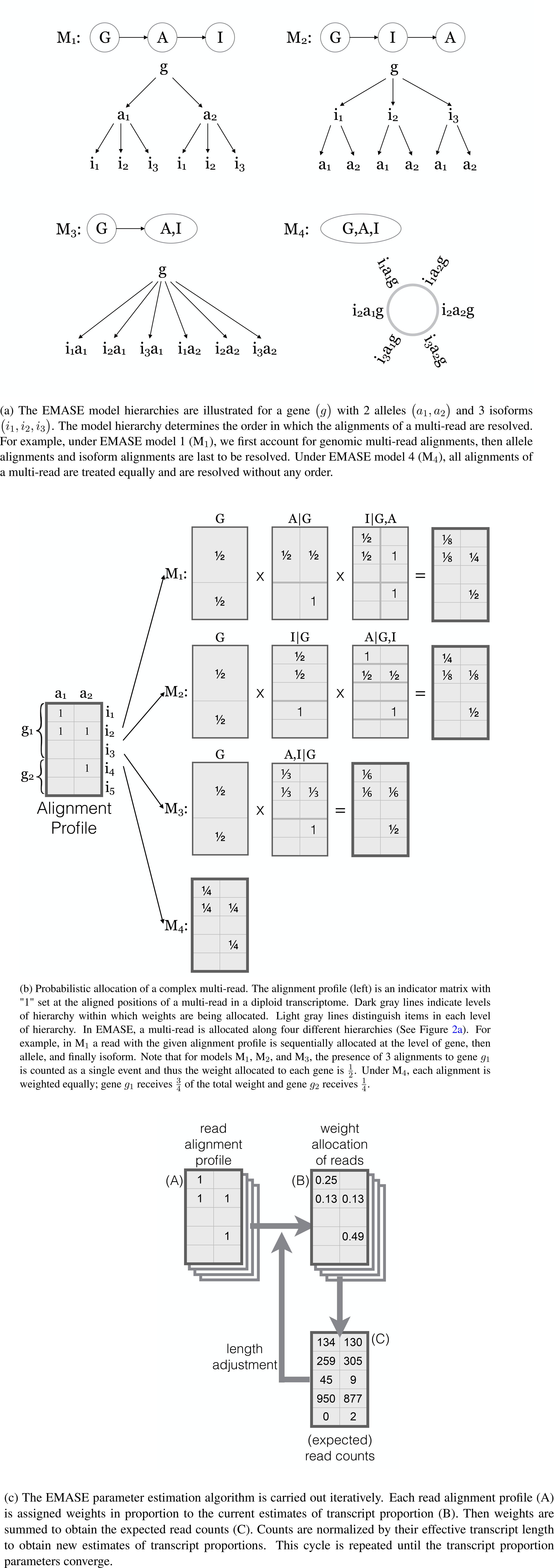
Hierarchical allocation of multi-reads

We implemented four EMASE models (M_1_, M_2_, M_3_, and M_4_) with distinct hierarchical structures (Figure 2a). Each model apportions a complex multi-read differently. Under M_1_, reads are apportioned among genes first, then between alleles, and then among isoforms. Under M_2_, reads are apportioned among genes first, then among isoforms, and then between alleles. Under M_3_ reads are apportioned among genes first, then among each isoform-allele combination which are treated equally. Model M_4_ assumes no hierarchy and multi-reads are apportioned equally among genes, isoforms, and alleles. M_4_ is implicitly the model used by other EM approaches. Under M_4_, the gene-level allocation of reads will depend on the number of isoforms that are represented in the transcriptome; genes with more isoforms will receive proportionately higher weights in the allocation of reads that are both genomic and isoform multi-reads. To see why this may be problematic, consider a situation where new isoforms of a gene are discovered and added to the transcriptome. With the new transcriptome definition, this gene will receive a larger share of the read allocation but the evidence that the read originated from this gene has not changed.

To illustrate how the four EMASE models allocate multi-reads, we constructed a hypothetical example of an alignment profile (Figure 2b). This is a complex multi-read at all three levels of the hierarchy. M_1_ first allocates equal weight to each gene; it then allocates weight between the two alleles of gene *g*_1_; lastly it allocates weights to isoforms within each gene and allele. Models M_2_ and M_3_ make similar allocations but in different orders resulting in different overall allocation of weights. We note that all three model M_1_, M_2_, and M_3_ given equal weight to each gene. In contrast, M_4_ will apportion weights equally to each alignment such that gene *g*_1_ receives 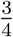 weight and gene *g*_2_ receives 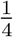 weight. In this example, we applied equally-weighted allocations to a single read. Next we describe an iterative algorithm for estimating the allocation parameters that uses data from all of the reads.

An EM algorithm is applied to obtain maximum likelihood parameter estimates for an EMASE model (Figure 2c). The EM algorithm for EMASE M_1_ begins with initial estimates of the relative expression of each gene(*θ_g_*), the allelic proportions for each gene (*ϕ_h|g_*), and the isoform proportions within each allele of each gene (*δ_i_*_|*g, h*_). Initial values can be equally weighted as in Figure 2b. The product of these parameters λ*_g_,_h, i_* = *θ_g_* · *ϕ_h_*|_*g*_ · *δ_i_*_|*g, h*_ represents the relative abundance of the transcript. In the E-step, current estimates of transcript abundance are used to apportion the unit count of a multi-read among the gene(s), isoform(s) and allele(s) to which it aligns. This process is repeated for each read and the weights are summed across all reads to obtain an expected read count for each transcript. In the M-step, the expected read counts are used to re-estimate the transcript abundance, incorporating an adjustment for the transcript length. The EM algorithm converges to yield maximum likelihood parameter estimates of transcript abundance and expected read counts. We note that expected read counts may not be integers due to the weighted allocation of multireads. Transcript abundance is a proportion among all transcripts and it is typically standardized to transcripts per million (TPM). Convergence of the EMASE fitting algorithm is declared when the sum of the absolute TPM changes by less than 1.0 on consecutive iterations. A detailed description of the EM algorithms is provided in Methods.

### Performance of EMASE on simulated data

We used simulations to evaluate the performance of EMASE models for estimating total and allele-specific expression and for comparison of EMASE to other approaches. We simulated 12 independent sets of 10 million 68 bp single-end reads using RSEM (Li and Dewey, 2011) version 1.3.0 with input parameters obtained by applying RSEM analysis to eight samples of F1 hybrid cross between mouse strains NOD/ShiLtJ (NOD) and PWK/PhJ (PWK). For the alignment phase of analysis, we generated NOD and PWK transcriptomes by incorporating known strain-specific SNPs and short indels into the reference transcriptome and combined these to form the diploid transcriptome of a NODxPWK F1 hybrid mouse using `g2gtools` (https://github.com/churchill-lab/g2gtools). We built the bowtie (Langmead *et al*, 2009) index using `rsem-prepare-reference` and aligned each of the simulated read sets to the diploid NODxPWK transcriptome using the `bowtie` aligner to generated BAM files for each of the 12 simulated data sets. Details of the simulations are provided in Methods.

We fit each of the four EMASE models with these BAM files and compared the accuracy of ASE and total expression estimation on the 12 simulated samples (Table 1). To determine the accuracy of ASE, we computed the proportion of genes or isoforms for which the absolute differences between estimated and true ASE is less than 0.1 (Table 1a). For gene-level estimates models M_2_ and M_4_ were equally best in performance and M_2_ was best for isoform-level ASE estimation. When we compared EMASE with the other EM methods, RSEM and kallisto, they all fell within a few percentage points of one another with kallisto having only marginally lower accuracy. The unique-reads method and WASP have substantially lower accuracy compared to EM methods. At the gene-level, WASP estimates of ASE fell within 10% of truth for fewer than half of genes. Isoform level estimates are not available with WASP.

To evaluate estimation of total gene expression among the EM based methods we computed the proportions of genes or isoforms for which the relative difference between estimated and true values was less than 10% (Table 1b). At both gene and isoform-level, M_4_ was most accurate based on number of genes that fell within 10% of the true value (Table 1b). All of the EM based methods have similar accuracy for total gene expression (~85%) and for total isoform expression (~40%). Kallisto has slightly lower accuracy and, as expected, total expression estimates for EMASE models M_1_, M_2_, and M_3_ are essentially identical.

**Table 1:** Accuracy of ASE and total gene expression estimation (See Figure S1 and S2).

|  | EMASE models |  |  |  | Other methods |  |  |  |
| --- | --- | --- | --- | --- | --- | --- | --- | --- |
|  | M<sub>1</sub> | M<sub>2</sub> | M<sub>3</sub> | M<sub>4</sub> | RSEM | kallisto | Unique | WASP |
| % of genes | 72.5±0.2 | **78.2**±0.3 | 77.9±0.3 | **78.2**±0.4 | 77.8±0.3 | 76.4±0.3 | 69.8±0.3 | 42.6±1.5 |
| % of isoforms | 56.9±0.2 | **60.9**±0.2 | 60.0±0.3 | 60.4±0.3 | 59.0±0.2 | 58.5±0.2 | 33.3±0.3 | N/A |

|  | EMASE models |  |  |  | Other EM methods |  |
| --- | --- | --- | --- | --- | --- | --- |
|  | M<sub>1</sub> | M<sub>2</sub> | M<sub>3</sub> | M<sub>4</sub> | RSEM | kallisto |
| % of genes | 86.8±0.2 | 86.8±0.2 | 86.8±0.2 | **88.7**±0.2 | 85.8±0.3 | 83.7±0.2 |
| % of isoforms | 38.3±0.2 | 38.3±0.2 | 38.3±0.2 | **39.1**±0.1 | 38.8±0.2 | 36.7±0.2 |

We carried out head-to-head comparisons between all pairs of models to determine the proportions of genes or isoform for which each method provided estimates of ASE or total expression that are closer to the simulated truth. For ASE, we report comparisons of EMASE model M_2_ against the other methods (Table 2a and Supplemental Figures S4 and S5). For total expression, we report comparisons between EMASE model M_2_ and the other EM methods (Table 2b and Supplemental Figures S6 and S7).

For ASE estimation, we considered the estimates obtained from two methods to be equivalent if the difference between the absolute deviations of estimated values from the simulated truth is 5% or less. In comparison to the other EMASE models, the M_2_ estimates are more often closer to the true values. The difference is most pronounced in comparison to M_1_ at the gene-level where 11.5% of genes were better estimated by M_2_ and only 3.3% were better estimated by M_1_. The differences among the EMASE model comparisons are less pronounced at the isoform level. In comparisons to the other estimation methods, M_2_ is consistently best and the next best performance is from RSEM followed by Unique and kallisto. The performance of WASP is an outlier — M_2_ provided substantially better estimates of ASE for 26.2% of all genes.

We compared EMASE model M_2_ estimates of total gene expression to the other EM based methods. Estimates were considered to be equivalent if they are within 5% relative difference. At the gene level, estimates from EMASE models M_1_, M_2_, and M_3_ are essentially identical and the minor differences (0.1%) can be attributed to convergence of the EM fitting algorithm. In comparison to model M_4_, we see that the M_2_ estimates are closer to truth for 1.4% of genes and M_4_ estimates are closer to truth for 5.5% of genes. Model M_2_ outperforms both RSEM and kallisto at the gene-level. At the isoform level, we see very similar performance among M_1_, M_2_, and M_3_. Model M_4_ is closer to truth than M_2_ for 12.7% of isoforms and RSEM also outperforms M_2_ in this comparison. We conclude that the best estimates of total gene expression are obtained using either EMASE model M_4_ or RSEM, but we note that all of the EM methods are performing within a few percentage points of one another. For both ASE and total expression, EMASE model M_2_ was overall best but not best in every comparison.

**Table 2:** Head-to-head model comparisons. See Figure S4, S6, S5, and S7.

| Comparing with $M_2$ | EMASE models | | | Other methods | | | |
| --- | --- | --- | --- | --- | --- | --- | --- |
| | $M_1$ | $M_3$ | $M_4$ | RSEM | kallisto | Unique | WASP |
| **% of Genes:** |  |  |  |  |  |  |  |
| EMASE $M_2$ is better | 11.5±0.2 | 1.1±0.1 | 2.3±0.2 | 3.7±0.2 | 7.1±0.3 | 6.2±0.1 | 26.2±0.8 |
| Column method is better | 3.3±0.1 | 0.6±0.1 | 2.0±0.1 | 2.9±0.1 | 4.4±0.2 | 2.6±0.1 | 5.1±0.2 |
| **% of Isoforms:** |  |  |  |  |  |  |  |
| EMASE $M_2$ is better | 16.9±0.2 | 18.2±0.2 | 18.7±0.2 | 19.8±0.3 | 22.2±0.2 | 13.6±0.2 | N/A |
| Column method is better | 11.0±0.2 | 12.2±0.2 | 13.1±0.2 | 13.2±0.2 | 14.0±0.2 | 5.6±0.1 | N/A |

| Comparing with $M_2$ | EMASE models | | | Other EM methods | |
| --- | --- | --- | --- | --- | --- |
| | $M_1$ | $M_3$ | $M_4$ | RSEM | kallisto |
| **% of Genes:** |  |  |  |  |  |
| EMASE $M_2$ is better | 0.1±0.0 | 0.1±0.0 | 1.4±0.1 | 7.6±0.3 | 11.9±0.2 |
| Column method is better | 0.1±0.0 | 0.1±0.0 | 5.5±0.1 | 5.7±0.2 | 5.5±0.1 |
| **% of Isoforms:** |  |  |  |  |  |
| EMASE $M_2$ is better | 4.2±0.1 | 6.9±0.2 | 10.4±0.2 | 21.6±0.3 | 27.4±0.3 |
| Column method is better | 4.7±0.1 | 6.4±0.2 | 12.7±0.2 | 25.6±0.2 | 25.0±0.3 |

It is often of interest to classify the ASE state of genes as monoallelic versus bi-allelic expression. In this simulation, we call a gene monoallelic if its estimated allele proportion is less than 2% or greater than 98%. It is also of interest to classify genes as expressed or not-expressed. We call a gene with expected read count less than 1.0 as not-expressed. Based on these classification rules, we compared the precision-recall of each model (Table 3).

The classification results for ASE are summarized in Table 3a. Precision — the proportion of monoallelic calls that are true — is around 50% for all of the ASE estimation methods with the exception of WASP for which only 16.4% of monoallelic calls were correct. Recall — the proportion of true monoallelic expressions that are called — is more variable across the ASE estimation methods with best performance reported for the unique-reads method followed by EMASE model M_4_, RSEM, and kallisto. At the isoform level, we can see a similar result. EMASE model M_1_ and M_2_ achieved the best precision, in excess of 60%, and the unique-reads method had the best recall of 93.7%, closely followed by EMASE model M_4_, RSEM, and kallisto.

Classification results for total expression are summarized in Table 3b. Precision and recall are consistently above 97% for each of the EM-based methods. Classification performance at the isoform-level is also consistently high across the EM-based estimation methods.

**Table 3:** Classification accuracy of monoallelic expression and gene expression (See Figure S1 and S2).

|  | EMASE models |  |  |  | Other methods |  |  |  |
| --- | --- | --- | --- | --- | --- | --- | --- | --- |
|  | M<sub>1</sub> | M<sub>2</sub> | M<sub>3</sub> | M<sub>4</sub> | RSEM | kallisto | Unique | WASP |
| **Genes:** |  |  |  |  |  |  |  |  |
| Precision | 0.515±0.019 | 0.541±0.017 | 0.540±0.017 | **0.550**±0.017 | 0.543±0.016 | 0.504±0.012 | 0.481±0.013 | 0.164±0.006 |
| Recall | 0.761±0.021 | 0.739±0.021 | 0.741±0.021 | 0.895±0.016 | 0.900±0.017 | 0.866±0.022 | **0.916**±0.015 | 0.541±0.038 |
| **Isoforms:** |  |  |  |  |  |  |  |  |
| Precision | 0.639±0.027 | **0.647**±0.018 | 0.549±0.016 | 0.555±0.014 | 0.539±0.014 | 0.511±0.016 | 0.508±0.015 | N/A |
| Recall | 0.456±0.017 | 0.571±0.012 | 0.817±0.012 | 0.929±0.012 | 0.929±0.012 | 0.899±0.013 | **0.937**±0.012 | N/A |

|  | EMASE models |  |  |  | Other EM methods |  |
| --- | --- | --- | --- | --- | --- | --- |
|  | M<sub>1</sub> | M<sub>2</sub> | M<sub>3</sub> | M<sub>4</sub> | RSEM | kallisto |
| **Genes:** |  |  |  |  |  |  |
| Precision | 0.977±0.001 | 0.977±0.001 | 0.977±0.001 | 0.972±0.001 | **0.982**±0.001 | 0.970±0.001 |
| Recall | 0.979±0.001 | 0.979±0.001 | 0.979±0.001 | **0.982**±0.001 | 0.980±0.001 | 0.974±0.001 |
| **Isoforms:** |  |  |  |  |  |  |
| Precision | 0.895±0.001 | 0.903±0.001 | 0.895±0.001 | 0.913±0.001 | **0.908**±0.001 | 0.905±0.001 |
| Recall | **0.852**±0.001 | 0.844±0.001 | 0.851±0.001 | 0.841±0.001 | 0.833±0.001 | 0.828±0.001 |

We examine the distribution of allele proportions in the simulated data (Figure 3a) and estimated allelic proportions by EMASE M_2_, RSEM, kallisto, WASP, and unique-reads methods (Figure 3). In total, ~22% and ~14% of reads are allelic unique at the gene and isoform level, respectively. EM-based methods (Figure 3b, 3c, and 3d) produced a smooth, bell-shaped distribution similar to the true distribution but with increased variation that reflects estimation error. WASP failed to estimate allele proportion in over 2,200 genes compared to the other methods. The overall distribution of ASE obtained from WASP was skewed toward NOD alleles, which are more similar to the mouse reference genome (Figure 3e). For these reasons, we conclude that postprocessing allele specificity after reference alignment is not fully correcting the alignment bias. The unique-reads method (Figure 3f) resulted in a symmetric distribution of allele proportions but with greater estimation error. We observed a 13% increase in variance at the gene-level and 39% increase at the isoform-level compared to EMASE model M_2_. Our implementation of the unique-reads method is unbiased due to the alignment to a customized transcriptome but estimates of ASE are more variable than EM and there are more monoallelic calls. EMASE model M_2_ reported the smallest number of false monoallelic expression calls: 168_±9_ versus 196_±13_ (EMASE M_4_), 203_±14_ (RSEM), 228_±12_ (kallisto), 264_±13_ (unique-reads method) or 735_±37_ (WASP) across 12 samples. See Table 3a for more detail.

**Figure 3:**
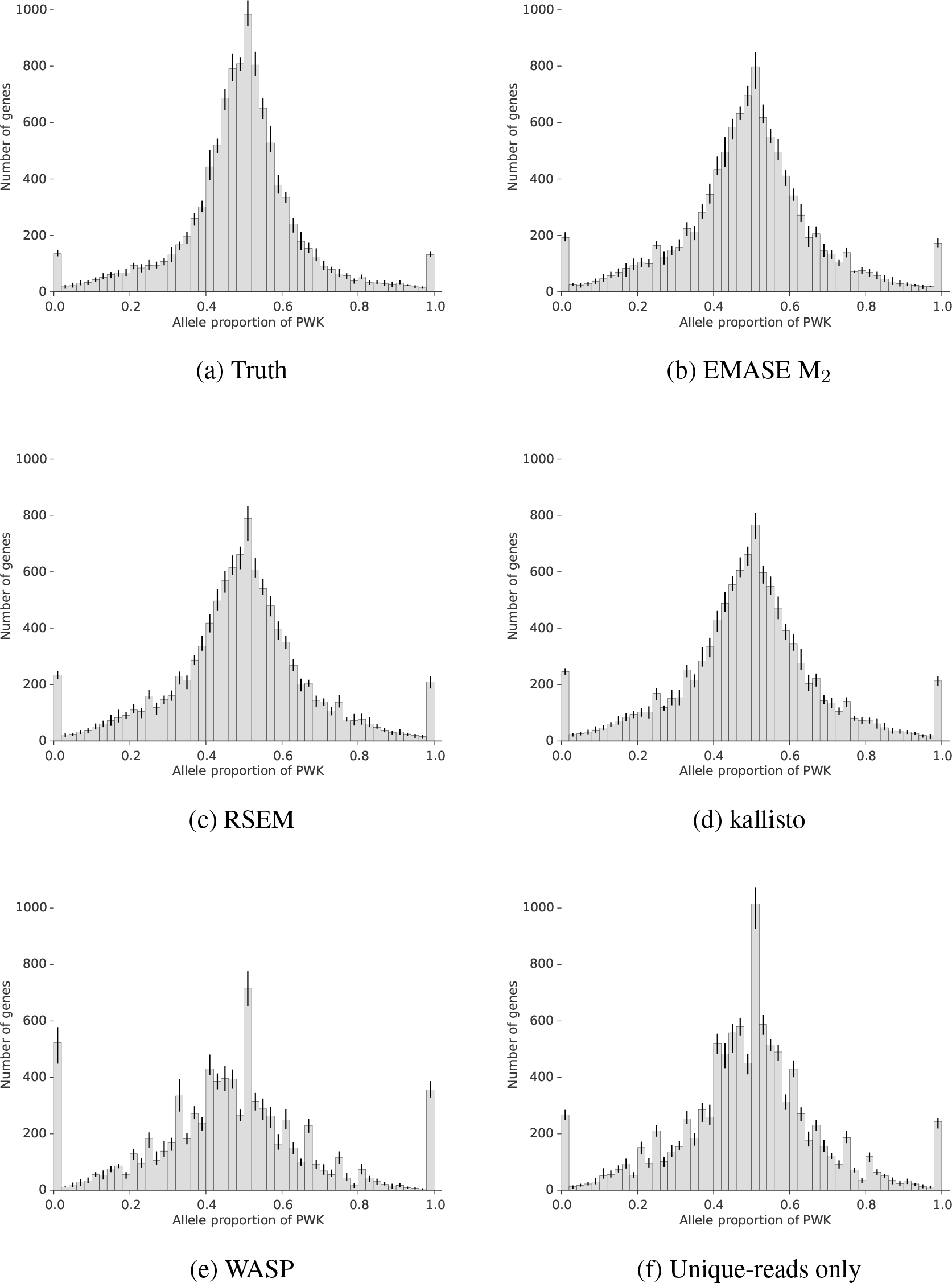
Estimated distribution of gene-level ASE from simulated data. Histograms show the frequency distribution across genes of the PWK allele proportions based on simulated truth (a), EMASE model M_2_ (b), RSEM (c), kallisto (d), WASP (e), and unique-reads only (f). The first and last bins in each histogram represent monoallelic expression defined as PWK allele proportion below 0.02 or above 0.98. Error bars are drawn between minimum and maximum values from 12 simulated data.

In summary, the EMASE models M_2_ and M_4_ estimates consistently provided the best or close to best estimates of both total expression and allelic proportion in our simulations. The same can be said for the RSEM estimates. Estimates obtained using kallisto were also consistently good but often not as accurate as the other EM methods — this may reflect some loss of information in the fast pseudo-alignment strategy. Among the non-EM methods, the unique-reads method (with alignment to the custom diploid transcriptome) provided consistent but less precise estimation. The WASP algorithm, which relies on a reference alignment strategy, performed poorly in all evaluations using simulated data.

### ASE in F1 hybrid data

We applied EMASE to RNA-Seq data from a reciprocal F1 hybrid cross between mouse strains NOD and PWK. There were 48 male mice in total with 24 mice from each direction of the cross (NOD×PWK and PWK×NOD). In order to evaluate the extent of ASE, we applied EMASE model M_2_ to estimate the PWK allele proportions for 9,102 informative autosomal genes (see Methods). This distribution of estimated ASE is symmetric (Figure 4a) indicating that there are no strain-specific biases. Monoallelic expression was observed for NOD alleles at 173 genes and for PWK alleles at 174 genes in the NOD×PWK samples. Monoallelic expression was observed for NOD alleles at 150 genes and for PWK alleles at 152 genes in the PWK×NOD samples. These are median values over the 24 samples in each cross direction. Numbers of monoallelic expressed genes varied from 115 to 454 in individual samples.

Male F1 mice from the two reciprocal crosses are hemizygous for the X chromosomes. We included both X chromosomes in our transcriptome definition in order to evaluate the misclassification rate of monoallelic expression. The majority of X chromosome genes (85% and 82% in NOD×PWK and PWK×NOD respectively) demonstrated monoallelic expression for the correct X chromosome (Figure 4a). Among the genes that show bi-allelic expression, 65% percent of the genes have fewer than 5 SNPs or indels to distinguish between alleles and others (35%) share sequence similarity and genomic multireads with autosomal genes.

**Figure 4:**
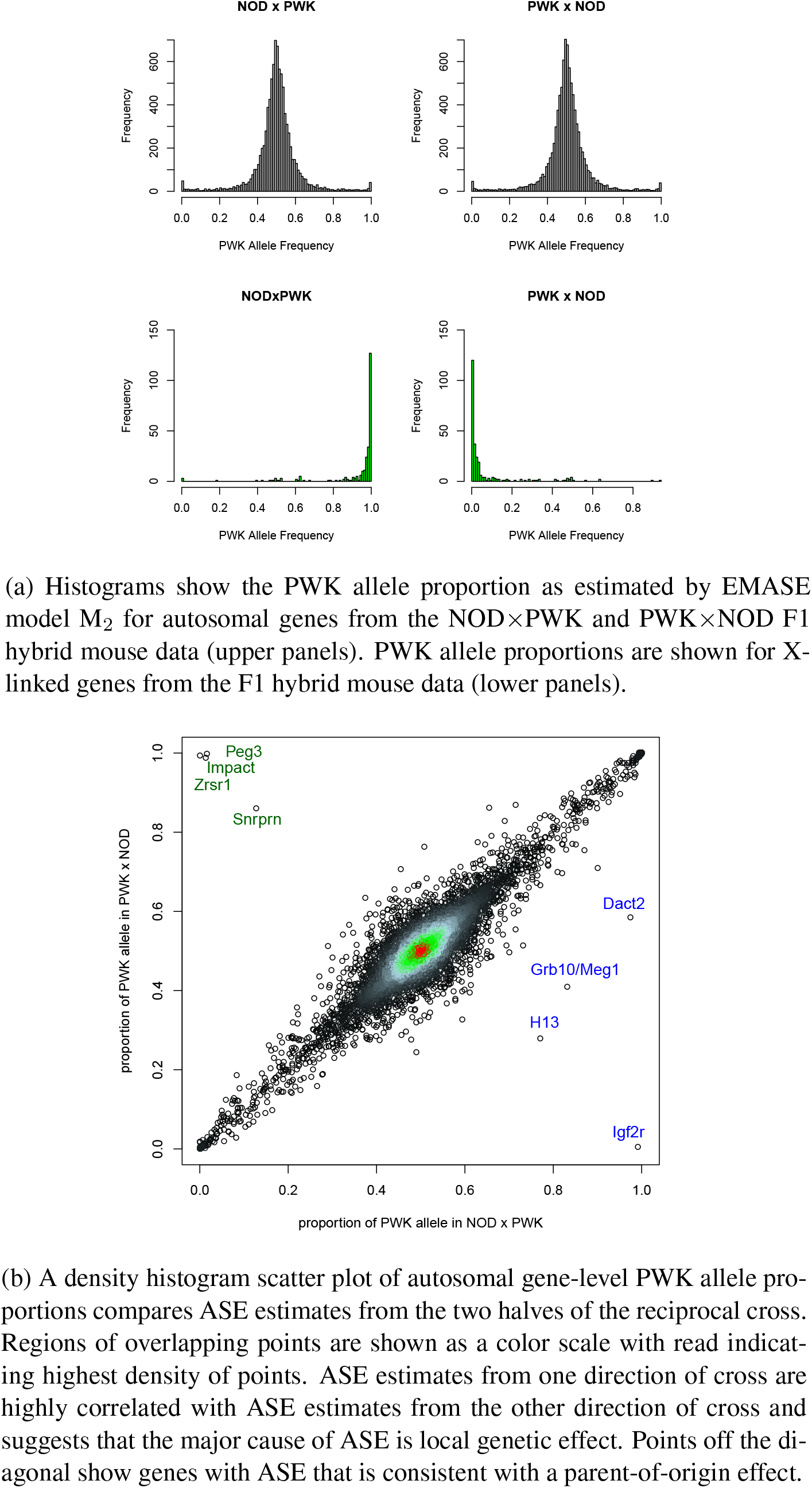
Allele-specific expression in a reciprocal F1 hybrid mouse population.

We evaluated the statistical significance of ASE using a beta-binomial test to account for overdispersion in (expected) read counts. We identified 4,216 genes (at FDR 5%) for NOD×PWK mice and 3,869 genes (at FDR 5%) for PWK×NOD mice with overlap of 3,084 genes (p<2.2*e*^-16^). This suggests that ASE is pervasive (affecting more then 35% of genes) and consistent across different groups of animals. A scatterplot of ASE estimates from each direction of cross reveals a striking level of concordance (*r*^2^ = 0.831) (Figure 4b) and suggests that ASE is continuously distributed with variable degrees of allelic imbalance across the genome.

We observed a handful of genes with a reversal in the PWK allele proportion between the two crosses, consistent with a parent-of-origin effect on ASE. In order to evaluate their significance we employed a logistic regression with a quasi-binomial likelihood (Agresti, 2002) and direction of cross as a predictor. We identified 70 genes with significant parent-of-origin effect, at 5% FDR (Figure S9). The strongest effects were restricted to genes that are already known to be imprinted, including *Igf2r, Peg3, Zrsr1, H13*, and *Impact*. We conclude that parent-of-origin effects are limited to a small number of well-characterized genes in adult mouse liver (Figure 4b). We also tested the effect of diet and age on ASE using the overdispersed logistic model and found 12 genes with significant diet effect and 112 genes with a significant age effect on ASE at 5% FDR threshold. These results suggest that allele-specificity is relatively insensitive to the diets and range of ages of mice in this study.

## Discussion

Until recently access to individual whole genome sequences has been out of reach for most organisms but sequencing of individual genomes of humans and model organisms is now proceeding rapidly. When individual genomes are not directly available, high-density genotyping arrays and variant databases can support imputation and phasing to obtain accurate approximations of individual diploid genomes. Hybrid mouse genomes, such as the NOD×PWK F1 animals used here, are straightforward to construct; they serve as a proof-of-principle for future applications of individually targeted RNA-Seq analysis. ASE estimation based on reference genome alignment suffers from bias even when secondary analyses are applied to account for misaligned reads. We recommend, whenever possible, to use an individually tailored diploid transcriptome incorporating known or imputed genetic variants as an alignment target for RNA-Seq analysis.

EMASE software works with BAM format files that can be produced by most short-read alignment software. It requires alignment to a collection of discrete sequence elements such as a transcriptome. The transcriptomes of human, mouse and other well-studied organisms are refined and well annotated. There is always room for improvement but for the present we rely on the reference transcriptome and adjust individual sequence elements to incorporate known or imputed SNPs and small indels. The impact of errors or individual variations in the set of the transcribed elements is not clear but will result in failure of some RNA-Seq reads to align to their correct origin or failure to align to any element. New alignment strategies that operate on whole genomes but are transcriptome-aware could help to address some of these concerns (Kim *et al*, 2015).

EMASE achieves up to 1,000 fold data reduction from BAM alignment format to the read alignment matrix. This reduction entails loss of information about the details of read alignments but this does not appear to impact the accuracy of estimation. Computing time for EMASE is substantially faster than RSEM but not as fast as *k*-mer methods such as kallisto. This suggests that detailed information about the aligned sequences is of limited value and that counting aligned reads is sufficient for accurate estimation.

The EMASE algorithm is readily adaptable to other contexts. All that is required is an alignment target composed of discrete sequence elements and a hierarchy. EMASE has been adapted to estimate allele-specific binding using ChIP-seq data (Baker *et al*, 2015) by defining sequence elements around DNA binding sites and applying a two-level hierarchy for sites and alleles. We have implemented an exon-junction version of EMASE as an alternative to the transcript isoform model presented here (Raghupathy et al., unpublished). EMASE has been adapted to analyze multiparent populations (http://churchill-lab.github.io/gbrs) with alleles assigned to eight (or any number of) haplotype classes (Chick *et al*, 2016). We anticipate the development of EMASE applications to allele-specific methylation, allele-specific RNA editing, and more.

Evaluation of RNA-seq analysis methods requires complex and realistic simulated data. Any simulation software makes assumptions that will affect the properties of the simulated data and the outcome of evaluations. After testing several simulation tools (Griebel *et al*, 2012; Frazee *et al*, 2015) including our own EMASE simulator, we decided to base our simulation studies on the RSEM simulator (Li and Dewey, 2011), which we found to be well documented and easy to implement. Moreover, we wished to avoid biasing our conclusions by evaluating the EMASE algorithm with data simulated from the EMASE model. We obtained input parameters for the simulations using values estimated from our F1 hybrid mouse data by RSEM. Thus both the input parameters and the simulated data were obtained from a non-hierarchical model that is most similar to EMASE model M_4_. It is difficult to assess the extent to which our simulations might have favored RSEM or EMASE model M_4_. RNA-Seq simulations can be sensitive to other choices such as whether and how to simulate poly-A tails and how to introduce sequencing errors. We made the choice to compare estimated read counts and allelic- or isoform-proportions of read counts as the outcome measure for comparison of different analysis tools. Alternative measures such as transcripts per million (TPM), are widely used but we found that each of the software tools has its own unique way of computing the transcript length adjustment for converting read counts to TPM. Read counts are the starting point for most normalization and downstream analysis methods, for example in voom (Law *et al*, 2014), edgeR (Robinson *et al*, 2010), and DESeq2 (Love *et al*, 2014), and they are consistently defined by the analysis methods considered here. Establishing standards for simulation-based evaluation of RNA-Seq analysis is an area in need of further attention.

EMASE estimates read counts at the level of transcriptome elements corresponding to individual isoforms and alleles of a gene. It can also aggregate counts at the gene-level and report average ASE across isoforms or isoform usage across alleles. If allelic proportions vary widely between isoforms, the gene-level average ASE may have little meaning. On the other hand when there is a dominant isoform or when ASE is consistent across isoforms, the gene-level summary will be more accurate due to the larger numbers of reads that are available to estimate ASE. Aggregate summaries of ASE are useful but should be viewed with caution. To obtain the best estimates of ASE, we recommend fitting model M_2_ and aggregating counts across isoforms — after checking that the isoform level estimates are not widely divergent.

EMASE explicitly models the different types of multi-reads and uses a hierarchical strategy to allocate weights. Our first implementation of EMASE was based on the hierarchy of model M_1_. It seemed logical because transcription acts first on an allele and splicing follows to produce the isoform. Yet model M_1_ consistently underperforms in comparison to other EMASE models. To understand the difference between models M_1_ and M_2_ in particular, we note that 86% of all reads are genomic unique, 50.6% of reads are isoform-unique, and 23.1% of reads are allele-unique. Thus we have more information to distinguish among isoforms than we have to distinguish among alleles. In addition, allelic-unique reads are typically defined by one or by a small number of SNPs; whereas isoform-unique reads can be distinguished across most or all of the nucleotides in the read. As a result we have more information to accurately allocate weights across isoforms compared to alleles. And we have the most information available to allocate multi-reads across genes. Our original motivation for constructing the EMASE hierarchy was based on the biology of transcription but a more pertinent consideration for determining the hierarchy is the information content of the data. We obtained the best results when we allocate at the most informative level first and the least informative level last. The differences in performance among the EMASE algorithms reflect this.

Ambiguity in read alignment presents a significant challenge for RNA-seq analysis. While it is tempting to discard or ignore multi-reads, this can lead to bias and reduced precision in estimation. Ambiguity in read alignment can be addressed by proportionately allocating counts using an EM algorithm, a general approach that outperforms methods that discard multi-reads. There are several EM algorithm implementations available, including EMASE, and they all perform well in head-to-head comparisons. EMASE resolves multi-reads by specifying a hierarchy among genes, isoforms, and alleles and we have found that the hierarchy of EMASE model M_2_ has generally the best performance. However, the differences reported in our evaluations are small and we would recommend the use of any these EM methods in practice.

## Methods

### Construction of the diploid transcriptome

We obtained NOD and PWK-specific SNPs and insertions and deletions (indels) of less than 100 bp from the Sanger Mouse Genomes Project SNP and indel Release Version 4 and mm10 genome (ftp://ftp-mouse.sanger.ac.uk/REL-1410-SNPs_Indels/). We used `g2gtools` (https://github.com/churchill-lab/g2gtools) to construct the strain-specific genome and strain-specific gene annotations using Ensembl release 75 *Mus_musculus.GRCm38.75.gtf* (ftp://ftp.ensembl.org/pub/release-75/gtf/mus_musculus) (Keane *et al*, 2011) and constructed strain-specific transcriptomes including all annotated isoforms. We merged the two transcriptomes into a single FASTA file and appended labels to track the strain-specific origin of each isoform.

### Read alignment and isoform abundance quantification

We built a bowtie index using `rsem-prepare-reference` from RSEM version 1.3.0 (Li and Dewey, 2011) with the NOD×PWK diploid transcriptome. We set the *polyA-length* parameter to attach a 67 bp-long poly-A tail to each transcript sequence element. We aligned our 68 bp singleend RNA-Seq reads to this diploid transcriptome using bowtie version 1.1.2 (Langmead *et al*, 2009) with parameters *‘–best’, ‘–strata’, ‘-a’, ‘-m100’* and *‘-v3’*. With these parameter settings, all read alignments with the best alignment score allowing up to 3 mismatches were retained for analysis. We used the same diploid transcriptome for building kallisto (version 0.43.0) pseudo-alignment index with default k-mer size of 31. We note that kallisto clips off the poly-A tail longer than 10 nt by default. We set the mean and standard deviation of fragment lengths to 250 and 25 respectively for both `rsem-calculate-expression` and kallisto quant.

### RNA-Seq simulations for benchmarking

We estimated isoform expression and allele proportion using `rsem-calculate-expression` on eight of our NOD×PWK samples (Sample IDs: 532702, 532704, 532706, 532708, 532710, 532714, 532716, and 532720) that contain ~40.4M aligned reads overall. We then simulated 12 independent sets of 10M reads using `rsem-simulate-reads` with the estimated parameters from the real data. For each simulation, we estimated expression at allele, isoform, and gene levels under all four EMASE models, RSEM, kallisto, and WASP. We started `rsem-calculate-expression` and EMASE from the same BAM files obtained using bowtie with the same index and parameters specified in the previous section.

### Animals and experimental design

We obtained male and female NOD/ShiLtJ (Stock No. 001976) and PWK/PhJ (Stock No. 003715) mice from The Jackson Laboratory (Bar Harbor, ME). Animals were received at 3 to 5 weeks of age, and given free access to food and water. We fed mice one of two diets from Harlan Teklad Custom Research Diets with matched ingredients with the exception of cholecalciferol. The standard diet contained 0.009 g/Kg and the enriched diet contained 0.050 g/Kg of cholecalciferol. NOD and PWK mice were mated using a reciprocal cross design (Figure S8) to obtain 24 male F1 progeny from each cross direction. We euthanized six mice from each cross-by-diet cohort at 15 weeks of age, and the remaining mice were euthanized at age 24 weeks. In total there are eight groups defined by the experimental factors (cross-direction, diet and age) and six biological replicates per group. The Animal Care and Use Committee at JAX approved the procedures in this protocol (AUS #99057).

### RNA sequencing

We collected liver tissue from the 48 F1 mice and stored it in RNAlater (Invitrogen) until processing. We isolated total RNA using the Trizol Plus RNA extraction kit (Life Technologies) with on-column DNase digestion, and purified mRNA from the total RNA using biotin tagged poly dT oligonucleotides and streptavidin beads. We fragmented mRNA by enzyme digestion, and generated double-stranded cDNA by random hexamer priming. We generated indexed mRNA-Seq libraries from 1_*μg*_ total RNA following the Illumina TruSeq unstranded protocol, and checked quality and quantity of the product using the Agilent Bioanalyzer and the Kapa Biosystems qPCR library quantitation method. We generated 68 bp single-end reads on the Illumina GAIIx sequencing platform. We assigned barcoded samples to sequencing lanes by randomization in a balanced design in which each lane included one sample from each of the treatment groups. We observed no significant differences between the technical replicates and for all subsequent analyses we used pooled data.

### Parameter estimation with EMASE

We first introduce some notations. Let *N_R_* be the number of RNA-Seq reads and let *r* = 1, …, *N_R_* be the index for reads. Let *g* = 1, …, *N_G_* be the index for genes, let *h* = 1,2 be the index for alleles, and let *i* = 1,… *N_I_*_|*g*_ be the index for isoforms of gene *g*. It will be convenient to denote the total number of isoforms in the transcriptome as 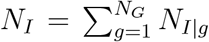 We denote the length of a transcriptome element as *l_ghi_* and assume the lengths are known constants with an appropriate adjustment for edge effects.

We denote true read origins as **R** = {*R_r_ : r* = 1, …, *N_R_*} a set of indicator matrices where *R_r_* is a matrix of size *N_I_* × 2 containing a single element with value one corresponding to the true origin of read r, and the rest zero. We denote read alignments as **A** = {*A_r_ : r* = 1,…, *N_R_}* a set of indicator matrices where A_*r*_ is a matrix with the same dimensions as R_*r*_ having one or more elements with value one corresponding to read alignments, and the rest zero. For uniquely aligned reads, the read origin and alignment matrices are assumed to be identical, *A_*r*_* = *R_*r*_*. For multiply aligned reads we assume that exactly one of the alignments correspond with the true origin of the read. Thus the element-wise product A_*r*_ × *R*_*r*_ = *R*_*r*_. The read alignment profile, *A*_*r*_, is observed data, but if r is a multi-read then *R*_*r*_ that represents the true origin of r is not directly observable. We also assume *R*_*r*_ given *r*’s alignment profile, *A*_*r*_, follows multinomial distribution of sample size 1 with probabilities, {*v_g,h,i|Ar_*} of length *N*_*I*_× 2. Our goal is to maximize the complete-data likelihood with respect to {*v_g,h,i|Ar_*}.

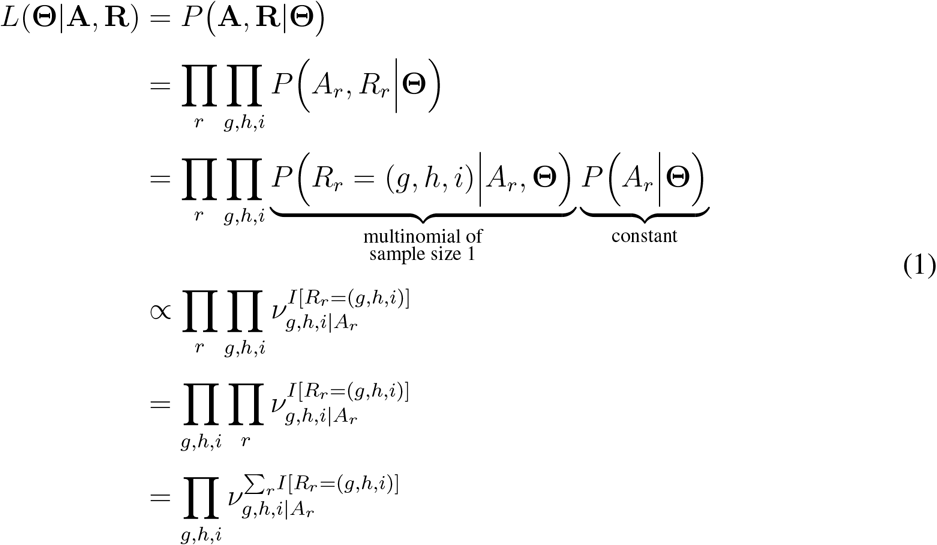

where *I[R_r_* = (*g, h, i*)], or *I_r_*(*g, h, i)* in short, is an indicator for whether the true origin of read *r* is isoform *i* of allele *h* of gene *g*. Note, given a transcriptome target, the alignment profile of a read is fixed, and therefore, *P(A*_*r*_ |Θ is constant. This equation shows Σ_*r*_*I*_*r*_ (*g, h, i)* is a sufficient statistic of `P(A, R`|Θ). Depending on the model of hierarchy, `I`_*r*_ (*g, h, i)* is defined in a different way and {*v_g,h,i|Ar_*} should be estimated accordingly. For example, for Model `M`_1_,

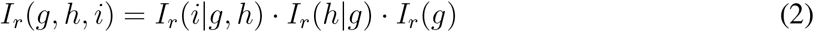

where

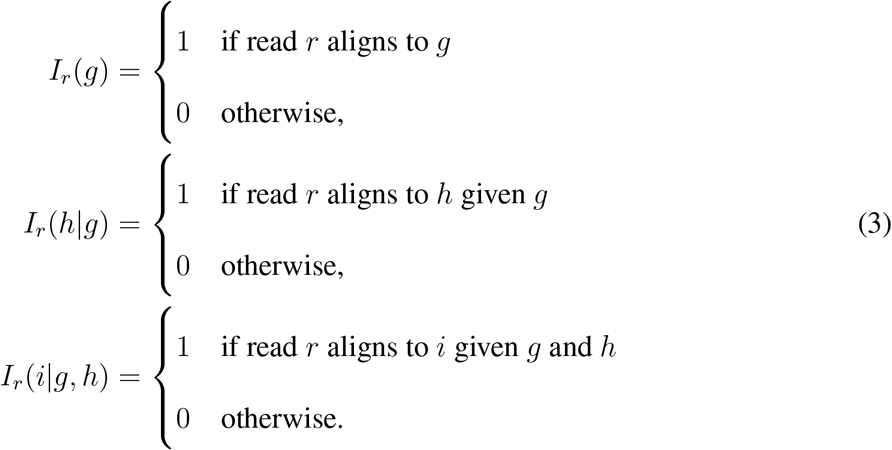

We also define additional parameters, *θ_g_*, *ϕ_h_*_|*g*_, and *δ_i_*_|*g,h*_, as the transcript proportion allocated to gene *g*, the transcript proportion from gene *g* allocated to allele *h*, and the transcript proportion from gene *g* and allele *h* allocated to isoform *i* respectively. Based upon these Model M_1_ indicators and parameters, we will describe EM-algorithm for M_1_. Analogous expression for M_2_, M_3_, and M_4_ are shown in the Appendix section. The complete-data likelihood belongs to the exponential family and its sufficient statistics are derived in Equation (1). The EM-algorithm is simple in that case as it reduces to updating the expectation of the sufficient statistic.

**E-step**: We compute *λ_g,h_,_i_*, the expectation on the sufficient statistic Σ_*r*_*I*_*r*_ (*g, h*, *i*) across all the possible combination of *(g, h, i)* given the observed data A and current estimates of the transcript proportion parameters Θ. *λ_g_,_h_,_i_* can be thought of as the expected read counts for each transcript. Precisely we compute *λ_g_,_h,i_* for EMASE model M_1_ using the following equation.

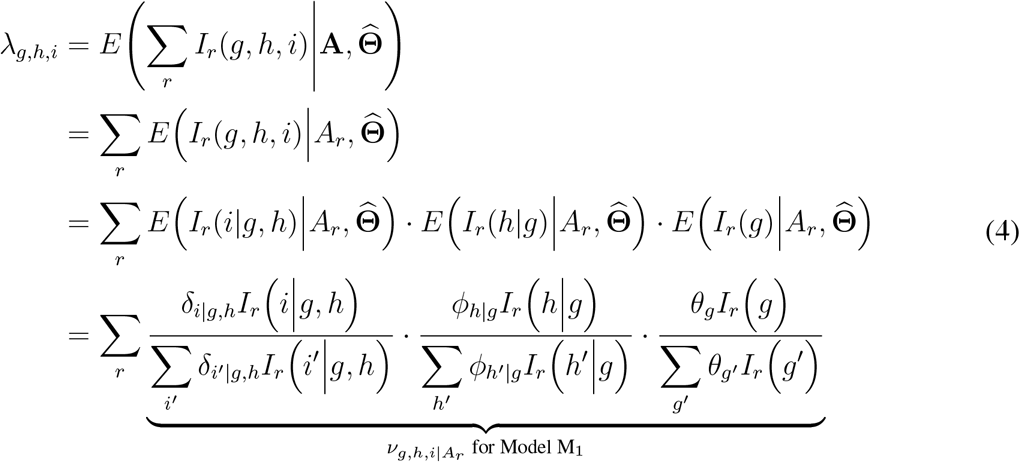

As previously mentioned, the quantities {*v*_*g h i|Ar*_} are the probabilities that govern the multinomial outcome of *R_r_* conditional on A_*r*_. They can also be thought of as weights that allocate read *r* proportionally across each of the elements to which it aligns.

**M-step**: We use the expected read counts, *λ_g_,_h_,_i_*, to estimate transcript proportion parameters Θ. For example, *δ_i_*_|g_,_h_, *ϕ_h_*_|*g*_, and *θ_g_* for M_1_ are obtained as

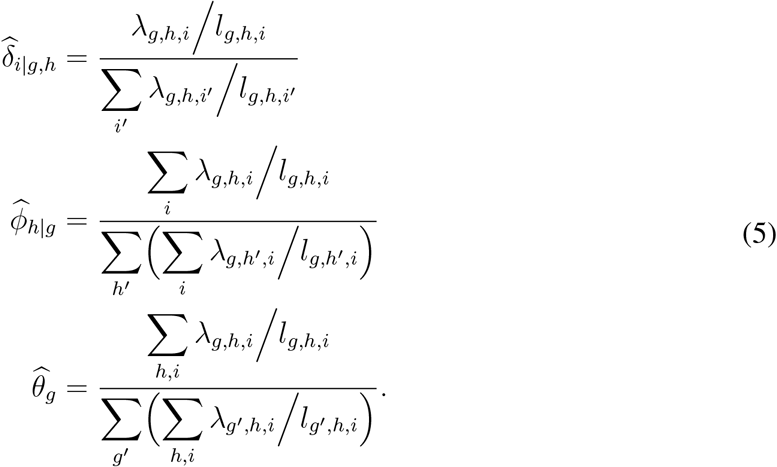

The length adjustment uses *l_g_,_h_,_i_*, the effective length of a transcript *i* of allele *h* of gene *g* which is defined to be [the length of the transcript] + [the length of the poly-A tail we added to the alignment target] - [the mean fragment length] + 1 (Nicolae *et al*, 2011; Li and Dewey, 2011).

### EMASE software

The EMASE software (DOI:10.5281/zenodo.291791) is implemented in Python and C++. It is available at https://github.com/churchill-lab/emase and is distributed under the GNU General Public License v3.0. We assume diploid genomes and haplotype-specific gene annotations are available or created using a package, e.g., `g2gtools` (DOI:10.5281/zenodo.292952) available at https://github.com/churchill-lab/g2gtools. These software tools take reference genome sequence file(s) in fasta format and variant call file(s) in vcf format and creates diploid pseudogenomes. `g2gtools` also lifts over reference gene annotation file (in GTF format) to a new pseudogenome coordinate system and extracts isoform (or genomic) sequences from the pseudogenome. EMASE command `create-hybrid` creates a diploid transcriptome by pooling two haploid transcriptome with allele identifiers added as suffixes and builds indices for read alignment. The reads can be aligned to the target sequences using any standard alignment software, such as `bowtie`. The command `bam-to-emase` converts the BAM format alignment file (Li *et al*, 2009) into an alignment incidence profile. The alignment profile file is a 3-dim sparse matrix of size *N_r_* × *N_I_* × 2 stored in PyTables HDF5 format (Alted *et al*, 2002-2014; The HDF Group, 2000-2010). The sparse matrix has value 1 if a read aligns to a specific isoform of an allele. The alignment profile file can be further reduced in size by collapsing together reads with identical alignment profiles as equivalent classes (Nicolae *et al*, 2011). The collapsed sparse matrix in HDF5 format reduces storage more than 1,000 fold compared to the original BAM file. For example, 40.4M 68-bp reads (See Methods) aligned to the diploid transcriptome results in a BAM file of size ~4.7GB, while the collapsed alignment profile file in HDF5 format is just about 3.1MB. The command run-emase executes the EM algorithm to quantify expression abundance at gene, allele, and isoform level. The user can optionally specify the convergence criteria and choose one of the four EMASE models (Figure 2a). According to our run test using the same 40.4M read data on a cluster compute node with Intel Xeon E5-2695 v3 @2.3GHz and 526GB RAM, EMASE C++ implementation ran 11.8 times faster than RSEM (5m59s versus 1h10m54s) using 16 cores in parallel, both starting from the same BAM file of ~270M alignments. For the same data and resource, kallisto took 1m41s which includes its own pseudoalignment process.

### Comparison methods

There are many software packages and workflows implemented for RNA-Seq analysis that provide estimates of ASE and total or isoform-level gene expression. These implementations differ in many small, and some important, details. For the purpose of evaluating EMASE we chose four comparative methods that reflect current practices in ASE estimation. Each method isolates one key feature in the analysis workflow. Using each of these methods, we estimate maternal (M) and paternal (P) allele-level read counts at the gene or isoform-level and estimate ASE as the proportion of maternal allelic read counts, *M*/(*M* + *P*). One method (WASP) does not produce isoform level estimates of ASE or total gene- and isoform-level expression. We excluded the unique-reads method for comparing total gene- or isoform-level expression because it loses too many reads for proper benchmarking. In order to focus just on the impact of discarding allelic multi-reads, we implemented our own version of this widely-used **unique-reads** approach to ASE estimation. We first aligned reads to the diploid transcriptome using the same BAM files as the EMASE and RSEM analyses. For gene-level ASE estimation, we discard the genomic multi-reads; and for isoform-level ASE estimation, we discard genomic and isoform multireads. Using the `count-alignments` feature of EMASE, we count reads that uniquely align to each allele at the gene- or isoform-level. **WASP** is an ASE estimation method that use a reference genome alignment. We aligned the reads to the mm10 reference genome using `TopHat` and used `WASP` software to re-assign the reads containing alternate alleles. The WASP corrected BAM files were used as input to `GATK`’s `ASEReadCounter` to get allele-level read counts at SNPs. The SNP level counts provide by WASP were averaged to get gene level ASE. We did not obtain estimates of isoform-level ASE with WASP. **RSEM** estimates ASE and total gene expression from aligned sequence reads. We used the same BAM format alignments for RSEM and EMASE analysis. RSEM retains multi-reads of all types and uses an EM algorithm to estimate expected counts. RSEM does not distinguish among allele, isoform and genomic multi-reads. **kallisto** estimates ASE and total gene expression based on a k-mer matching algorithms that generates a pseudo-alignment to RNA-Seq reads. It uses an EM algorithm to resolve multi-reads and, like RSEM, treats all types of multireads equally. We provide the NODxPWK diploid transcriptome to kallisto and obtain estimates of ASE and total expression at the gene- and isoform-level.

### Statistical significance of ASE

We used a Beta-Binomial model of individual sample allele-specific read counts across 48 samples to evaluate the statistical significance of ASE. The Beta-Binomial model allows sample-to-sample variation in ASE and has been shown to accurately model overdispersion in RNA-Seq data (Pickrell *et al*, 2010). If X is the expected read count from the PWK allele and *n* is the total read count, we assume that X is drawn from a binomial distribution with parameters *n* and *p*, *X* ~ *Bin(n,p)*, where *p* is a random variable drawn from a beta distribution with parameters *α* and *β*, *p* ~ *Beta(α, β*). We tested the null hypothesis H_0_ : *α = β*, which corresponds to values of p that are on average 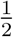, against the alternative hypothesis H_0_ : *α ≠ β* using a likelihood ratio test. We evaluated the significance of ASE separately for each direction of cross, treating each of the 24 samples from one direction as independent replicates and made no adjustment for diet or age at this stage.

In addition, we tested for effects of direction-of-cross, diet, and age on ASE using an additive logistic regression model (Cross+Diet+Age) with a quasi-binomial link function to account for overdispersion. We evaluated the significance of each main effect by dropping the corresponding term from the additive model and computing the likelihood ratio (chi-square) statistic for the reduced model.

### Software and Data Availability

The EMASE software is freely available under GNU license at https://github.com/churchill-lab/emase. Raw RNA-Seq data files in fastq format and processed allele-level expression estimates are archived at GEO under the accession #xxxxxx. Simulation data are available at ftp://churchill-lab.jax.org/pub/software/EMASE/.

## Author’s contributions

NR, KC, SM, and GA conceived the algorithm and KC, MV, GB, and KS implemented the methods. RK and FP provided data and interpretation of findings. KC and NR analyzed data. KC, NR and GAC wrote the manuscript.

## Appendix A: EMASE Models

As described in previous sections, the complete-data likelihood is

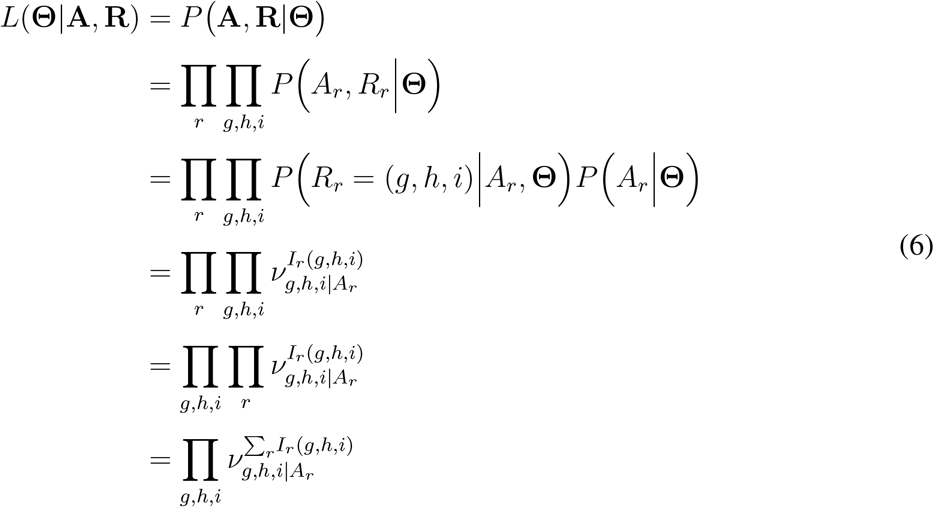

*P*(A, R|Θ) is the sum of multinomial random variables {R_r_}, each having sample size of 1 with the probabilities {*v_g h_*_*i*|*Ar*_}. The probability *v_g h i_*_|*Ar*_ is a function of A_r_ and *λ_g_,_h_,_i_*, the alignment profile of read *r* and the proportion of reads that originate from isoform *i* of allele *h* of gene g respectively. Depending on our models of hierarchy, *I_r_* (*g, h, i*) is defined in the following ways,

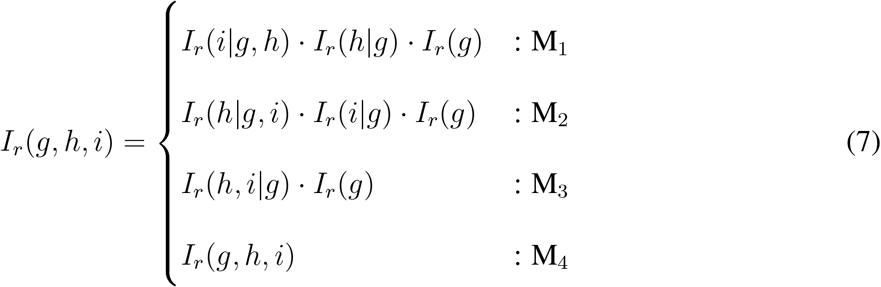

and *v_g h i_*_|*Ar*_ should be estimated along this structure. No nested structure is assumed in Model 4.

In E-step, we compute the expectation on the sufficient statistic Σ_*r*_*I*_*r*_ (*g, h, i*) given the observed data A and the current estimates of the model parameters Θ. Similarly to *θ*_*g*_, *ϕ_h_*_|*g*_, and *δ_i_*_|g_,_h_ for Model 1, we define *ϕ_h_*_|*g*_,_i_ and *δ_i_*_|*g*_ to be the transcript proportion that originate from allele h given isoform *i* of gene *g* and the transcript proportion from isoform *i* given gene *g* respectively for Model 2. For Model 3, we also define *δ_h,i_*_|*g*_ to be the transcript proportion from isoform i of allele h given gene g, and, for Model 4, *δ_g_,_h_*,_*i*_ to be the transcript proportion from isoform *i* of allele *h* of gene *g*.

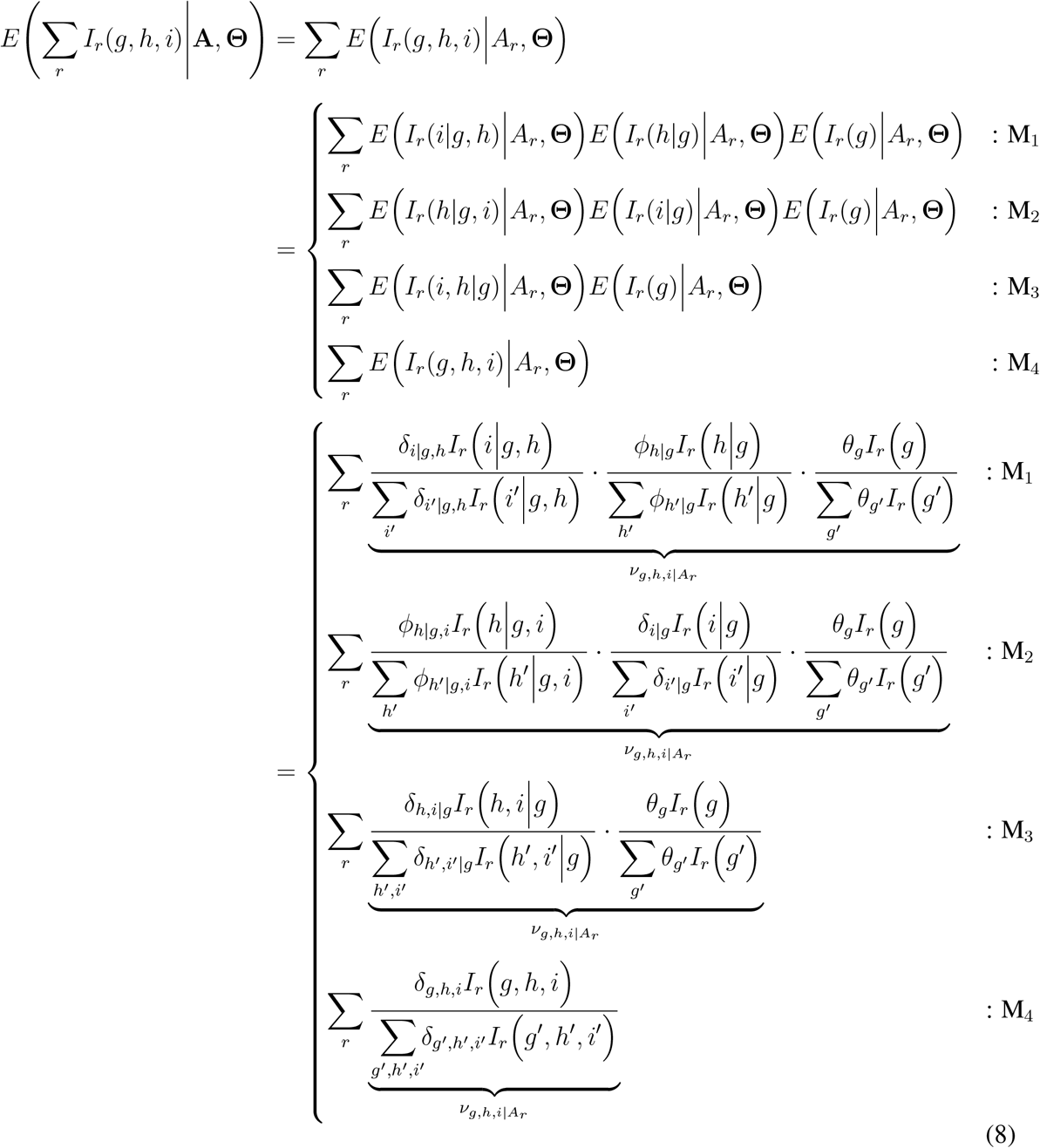

In M-step, we update the maximum likelihood estimator (MLE) of *δ, ϕ*, and *θ* in each model. Suppose λ_*g,h,i*_ denotes the proportion of reads sampled from transcript *i* of allele *h* of gene *g*, then, for all models,

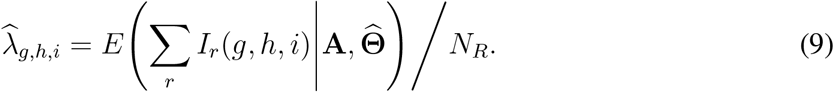

Since *δ*, *ϕ*, *θ* are all proportions with respect to the transcript count (or equivalently coverage depth),

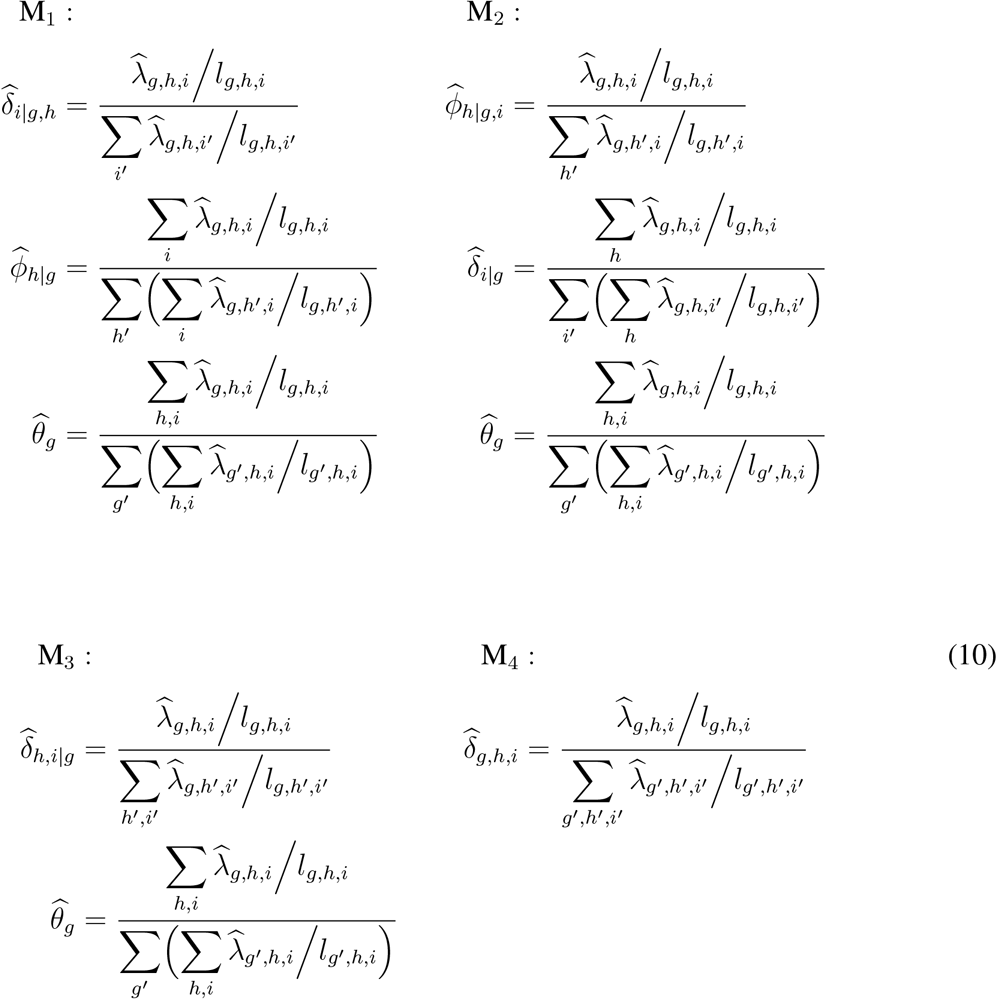

where *l_g,h,i_* is the effective length of isoform *i* from allele *h* of gene *g*: (the length of the isoform) + (the length of poly-A tail we add to the target sequences) - (the RNA-Seq read length) + 1.

**Figure S1:**
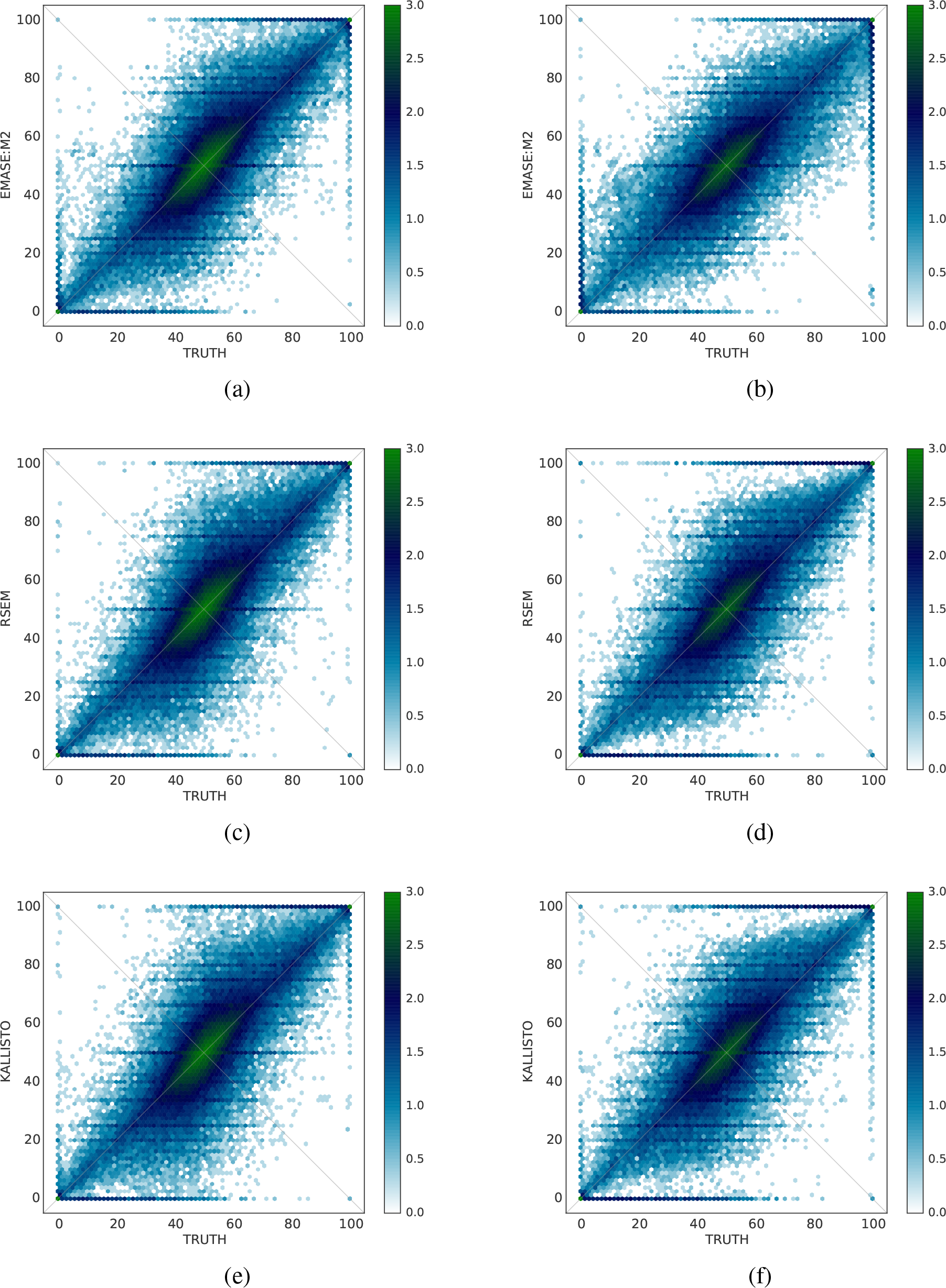
Performance of EMASE model M_2_, RSEM, and kallisto on ASE at the gene level (a)(c)(e) and at the isoform level (b)(d)(f)

**Figure S2:**
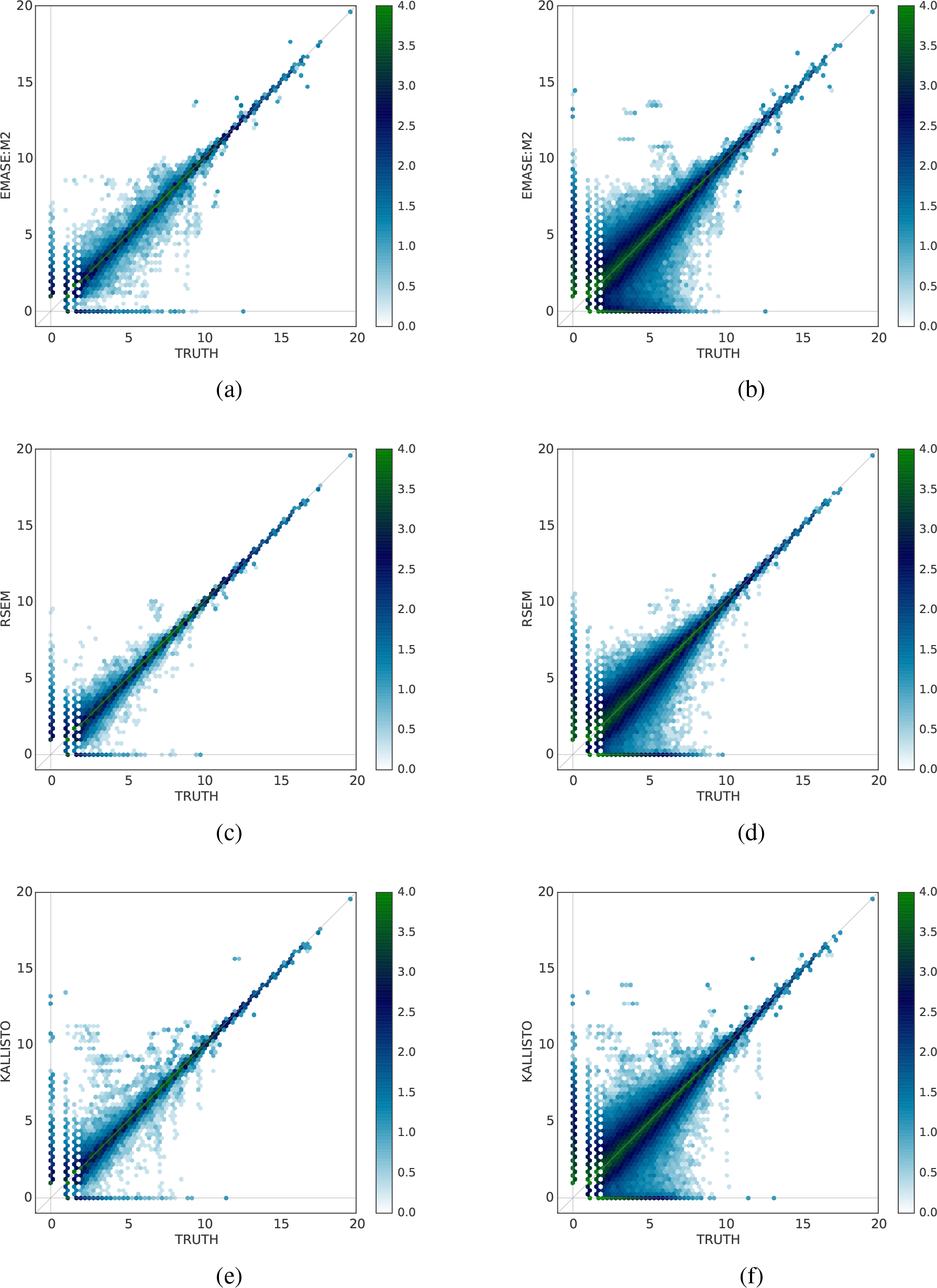
Performance of EMASE model M_2_, RSEM, and kallisto on total gene expression (a)(c)(e) and total isoform expression (b)(d)(f)

**Figure S3:**
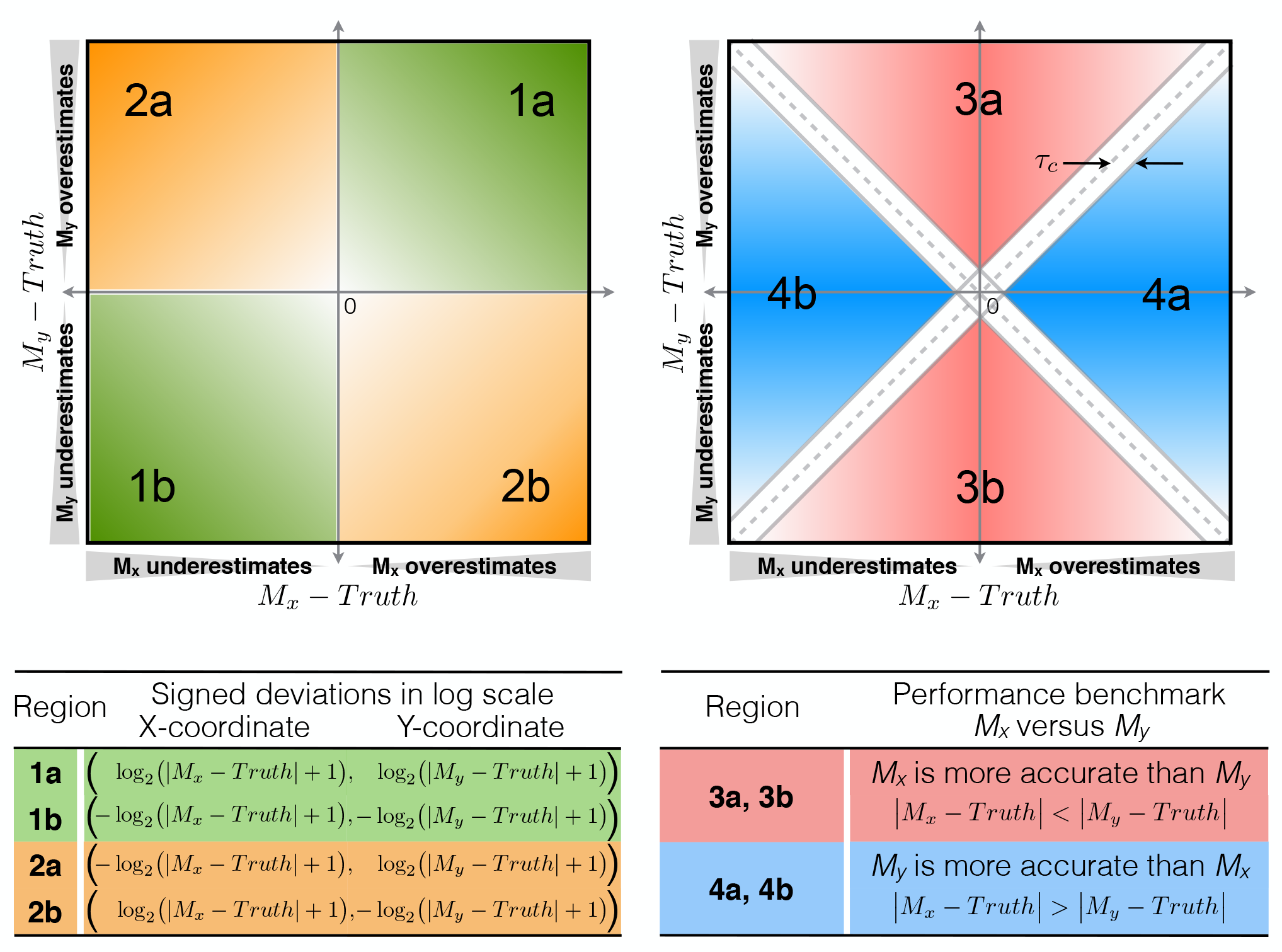
Signed deviation plot for head-to-head comparison of methods for ASE and total expression estimation methods. This figure provides a guide for interpreting the supplemental figures S4 through S7, which provide details behind the comparison summaries in Table 2. The x- and y-axes represent deviation of the estimated values from the true simulated values. Therefore, each gene or isoform is represented as a point at coordinate (*M*_*x*_ — *Truth, M_y_ — Truth*) where *M_x_* and M_*y*_ denote estimated values. The range of total expression estimates is wide and we converted them to log scale by taking logarithm on the absolute value of the differences after adding 1.0 and we keep the sign of the deviations. Thus negative values on each axis denote underestimation and positive values indicate overestimation relative to truth. On either scale, regular or log scale, a point nearer to y-axis (quadrants 3a or 3b) indicates higher accuracy of the *M_x_* estimates and a point nearer to the x-axis (quadrants 4a or 4b) indicates higher accuracy of *M_y_* estimates. Dotted lines are demarcate where the estimates of *M_x_* and *M*_*y*_ have essentially the same deviation from the truth. The proportions reported in Table 2 are obtained by comparing the count of items in 3a,b to those in 4a,b relative to the total numbers of genes or isoforms. This configuration is applied to plots in Figures S4, S5, S6, and S7 and log-scale is used for Figures S6 and S7.

**Figure S4:**
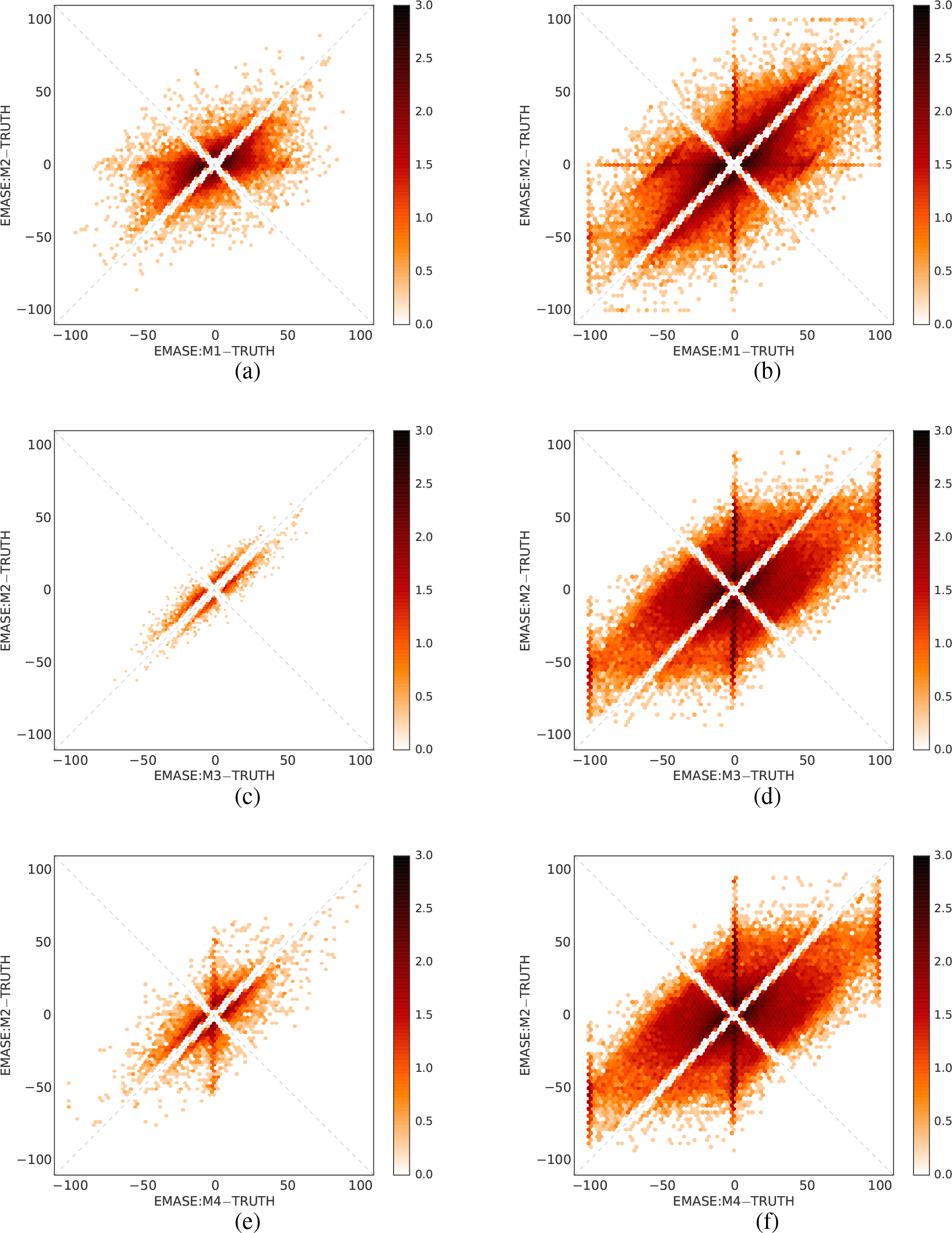
Head-to-head comparison between EMASE Models on ASE. Since M_2_ performed best overall, we show it compared with the other models with respect to deviations from the true proportions (%) of the PWK allele expression at the gene-level (a)(c)(e) and at the isoform-level (b)(d)(f). Frequency values in each hexbin cells in these plots are average across twelve samples. These frequencies are also rescaled in log_10_ space in order to visualize highly dispersed values.

**Figure S5:**
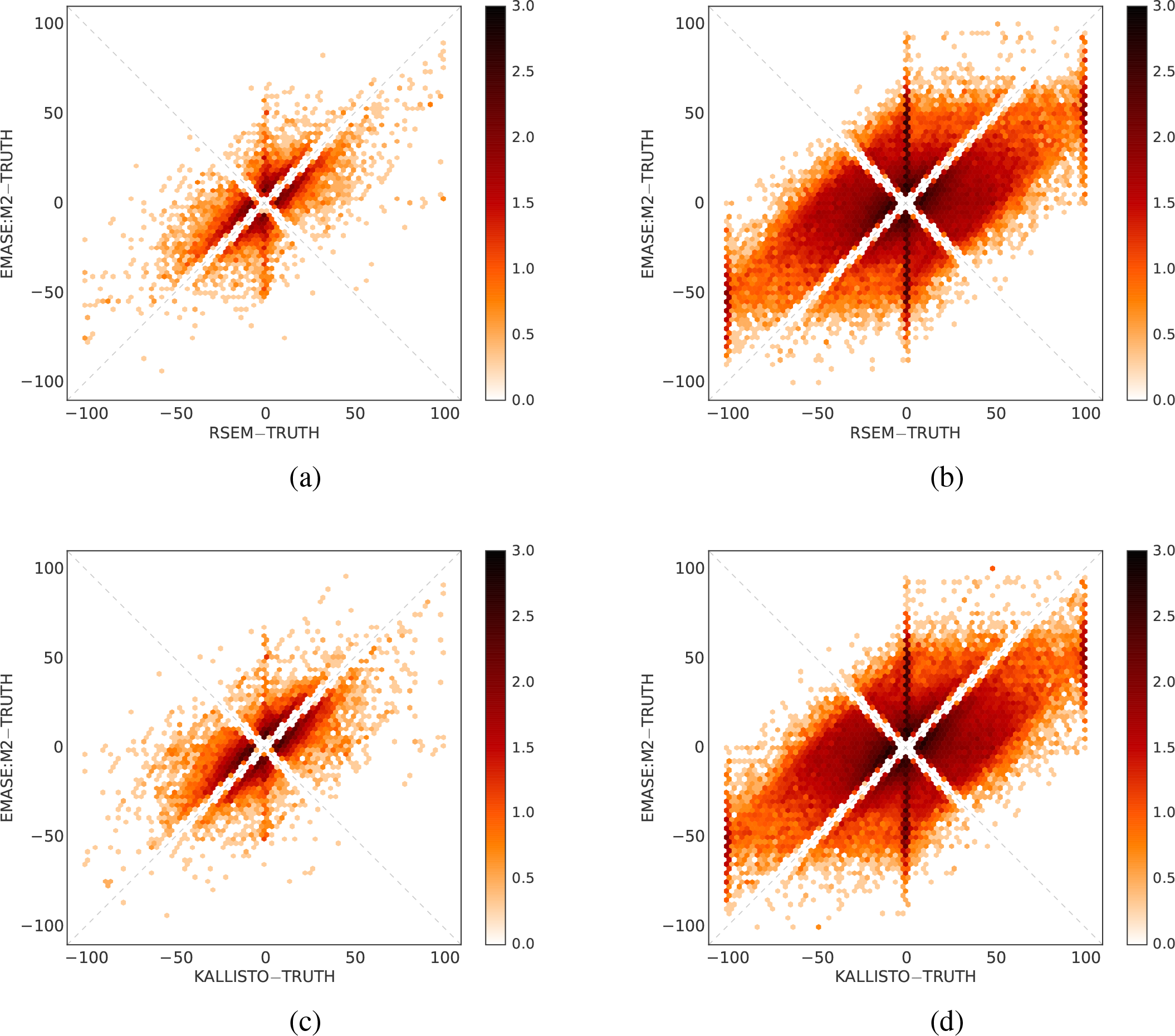
Head-to-head comparison between EMASE and other EM-based method, RSEM (a)(b) and kallisto (c)(d), on ASE. Comparisons are performed at the gene-level (a)(c) and at the isoform-level (b)(d). Frequencies in 2-dim histogram (or hexbin) are in log_10_ scale.

**Figure S6:**
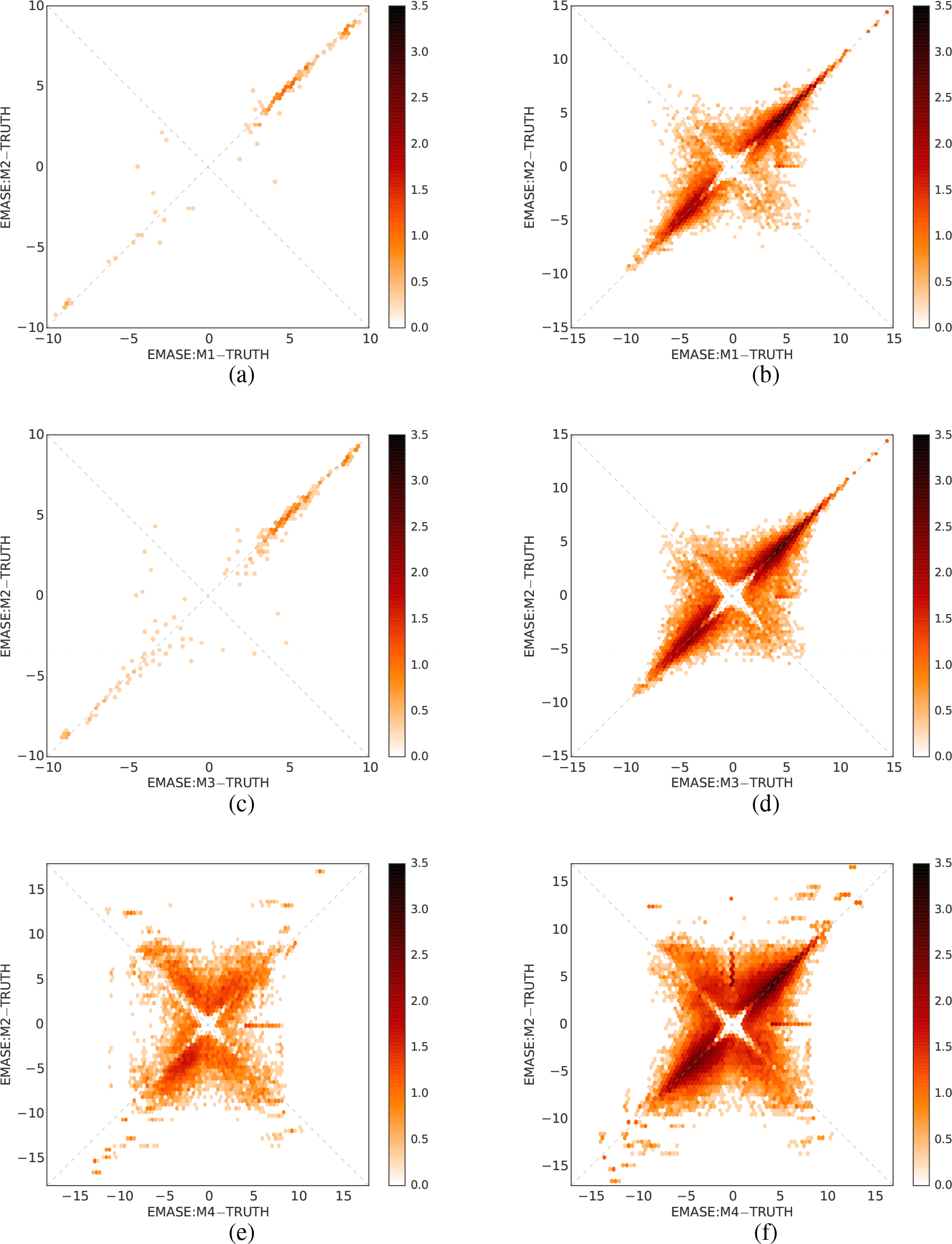
Head-to-head comparison between EMASE Models with respect to total gene expression (a)(c)(e) and total isoform expression (b)(d)(f). Again, we show how M_2_ performs compared with other models. Plots are in log scale (See Figure S3 for how to transform the signed deviation plot to log scale). Frequency values in each hexbin cells in these plots are average across twelve samples. These frequencies are also rescaled in log_10_ space in order to visualize highly dispersed values.

**Figure S7:**
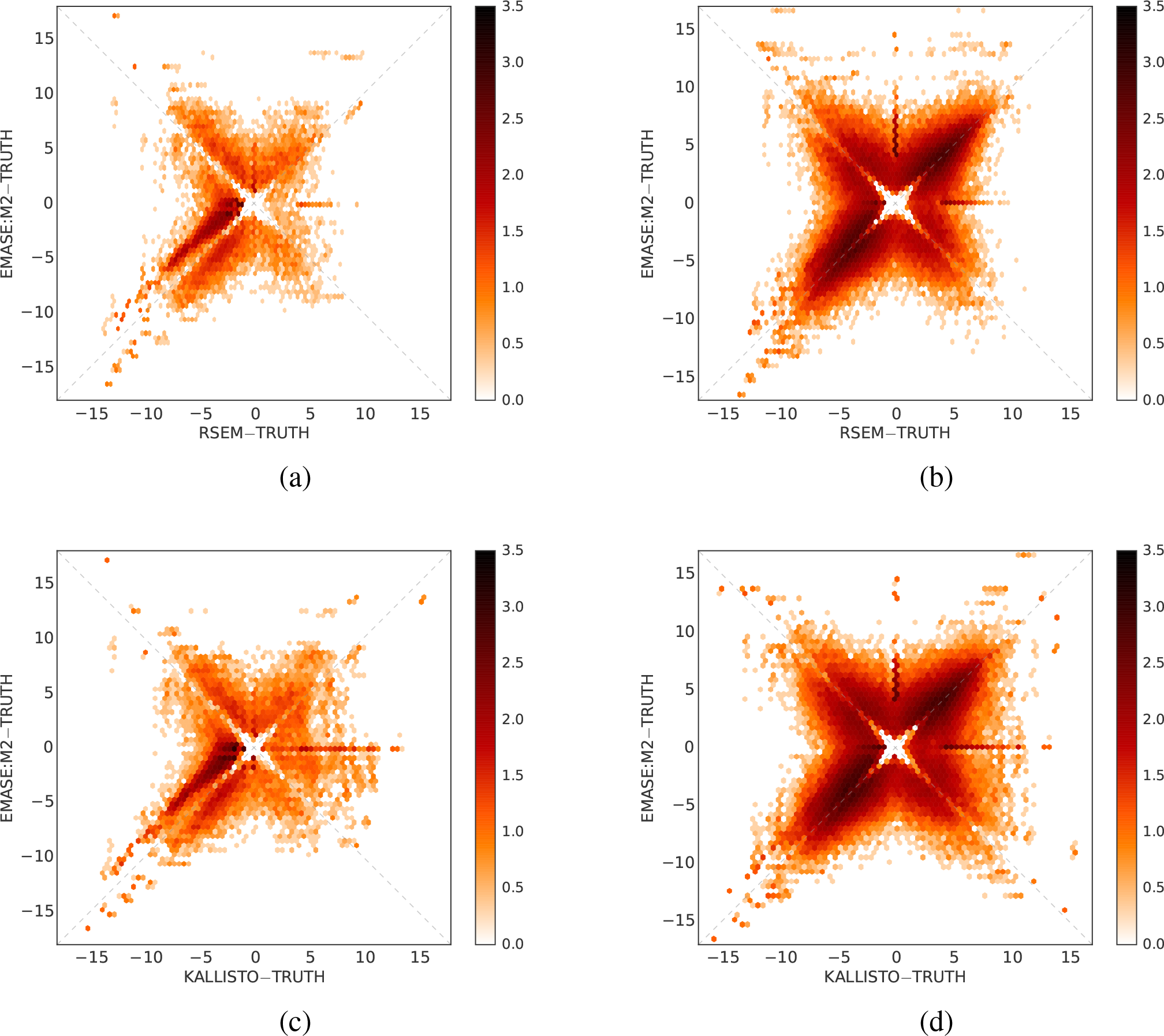
Head-to-head comparison between EMASE and other EM-based methods, RSEM (a)(b) and kallisto (c)(d), on total expression. Comparisons are performed at the gene level (a)(c) and at the isoform level (b)(d). Frequencies in 2-dim histogram (or hexbin) are in log10 scale. A darker area at the lower left region in the left quadrant of (a) and (c) indicate that EM methods generally underestimate expression but EMASE does it in a lesser degree compared to RSEM or Kallisto.

**Figure S8:**
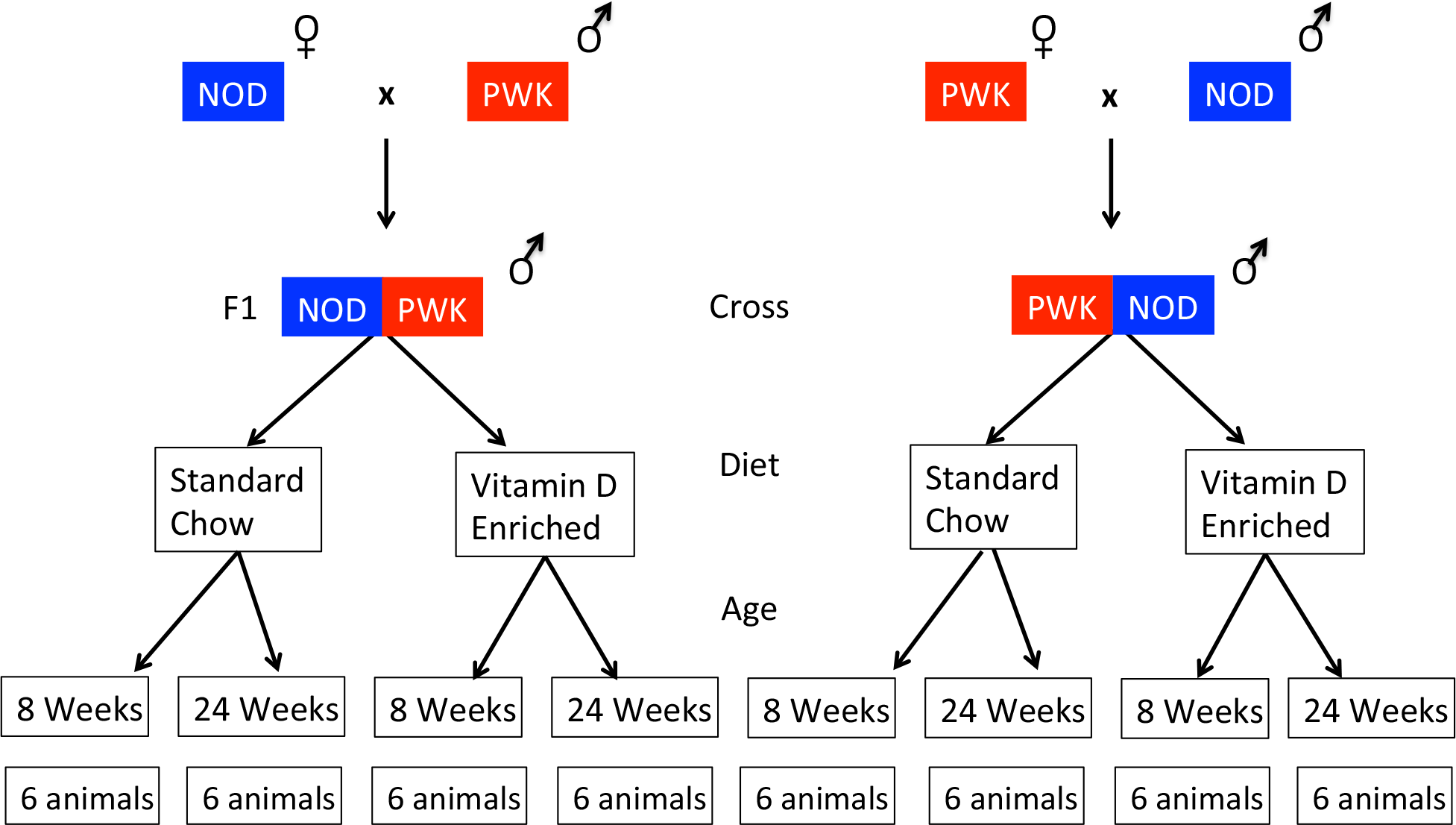
Experimental design of the Reciprocal F1 hybrid cross.

**Figure S9:**
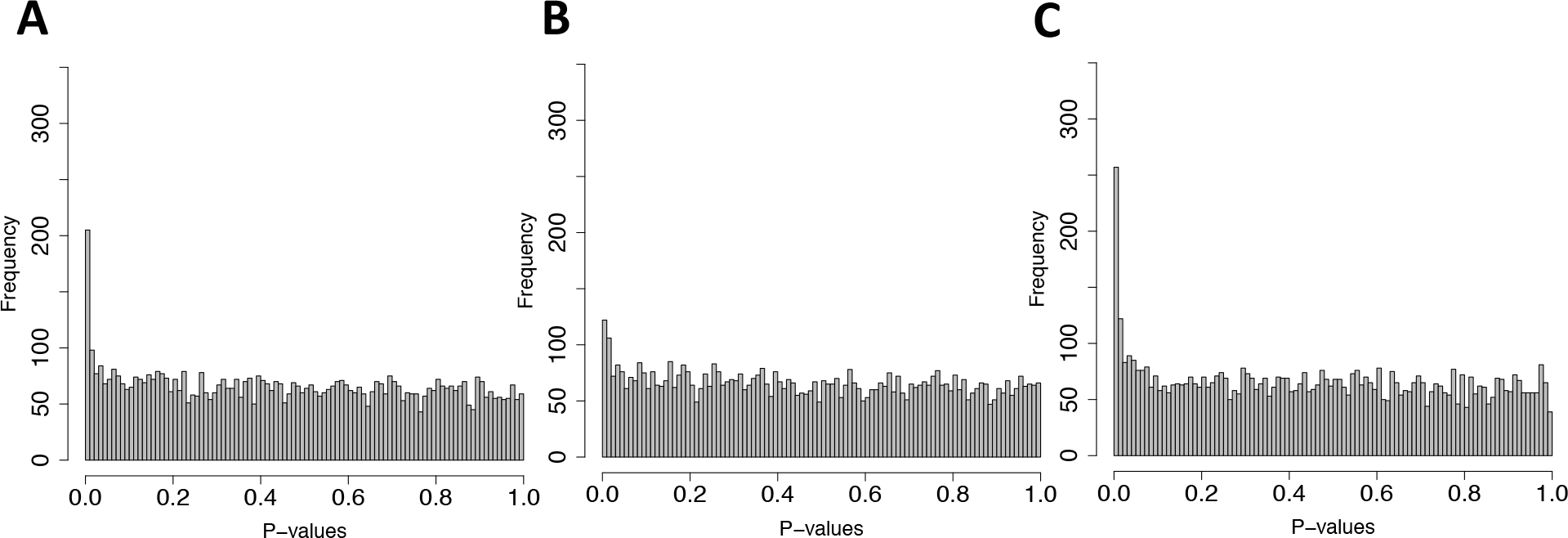
Effects of direction-of-cross, diet and age on ASE. P-value histograms for the effect of (A) direction of cross, (B) diet, and (C) age on ASE. This shows that direction of cross has small effect on ASE, diet has almost no effect on ASE and age has decent effect on ASE.

